# Morpho-molecular diversity and evolutionary analyses suggest hidden life styles in Spumellaria (Radiolaria)

**DOI:** 10.1101/2020.06.29.176917

**Authors:** Miguel M. Sandin, Tristan Biard, Sarah Romac, Luis O’Dogherty, Noritoshi Suzuki, Fabrice Not

## Abstract

Spumellaria (Radiolaria, Rhizaria) are holoplanktonic amoeboid protists, ubiquitous and abundant in the global ocean. Their silicified skeleton preserves very well in sediments displaying an excellent fossil record, from the early middle Cambrian (ca. 509-521 Ma), extremely valuable for paleo-environmental reconstruction studies. Spumellaria are tedious to maintain in laboratory conditions preventing an accurate perception of their extant diversity and ecology in today’s oceans, most of which being inferred from sediment records. This study represents an integrated and comprehensive classification of Spumellaria based on the combination of ribosomal taxonomic marker genes (rDNA) and morphological characteristics. In contrast to established taxonomic knowledge, we demonstrate that symmetry of the skeleton takes more importance than internal structures at high taxonomic rank classification. Such reconsideration allows gathering different morphologies with concentric structure and a spherical or radial symmetry believed to belong to other Radiolaria orders from the fossil record, as for some Entactinaria families. Results obtained in this study suggest the existence of new Spumellaria diversity at early diverging positions, in which a non-bearing skeleton organism lives within shelled ones. Using fossil calibrated molecular clock we estimated the origin of Spumellaria in the middle Cambrian (ca. 515 Ma), in agreement with the appearance of the first radiolarian representatives in the fossil record. The morpho-molecular and evolutionary framework established herein allows a direct connection between living specimens and fossil morphologies from the Cambrian, bringing both a standpoint for future molecular environmental surveys and a better understanding for paleo-environmental reconstruction studies.

## Introduction

Radiolaria are holoplanktonic amoeboid protists belonging to the Rhizaria lineage, one of the main branches of the eukaryotic tree of life (Burki and Keeling, 2014). Along with Foraminifera they compose the phylum Retaria, characterized by cytoplasmic prolongations and a skeleton composed of different minerals (Adl et al., 2019). Spumellaria represents an important order within the Radiolaria (Suzuki and Not, 2015), extensively studied across the fossil record thanks to its opaline silica skeleton (De Wever et al., 2001). Along with that of other radiolarians, spumellarian fossil record dates back to the early middle Cambrian (ca. 509-521 Ma; Suzuki and Oba, 2015; Aitchison et al., 2017; Zhang & Feng, 2019) constituting an important tool for paleo-environmental reconstructions analysis (e.g., Abelmann and Nimmergut, 2005). In contemporary oceans, molecular-based metabarcoding surveys performed at global scale have shown Radiolaria contribute significantly to plankton communities (de Vargas et al., 2015; Pernice et al., 2016) and silica biogeochemical cycle (Llopis-Monferrer et al., 2020), although very little is known about their ecology and diversity. Living Spumellaria, as other Radiolaria, are characterized by a complex protoplasmic meshwork of pseudopodia extending radially from their skeleton (Suzuki and Aita, 2011). With their pseudopodia, they actively capture prey, such as copepod nauplii or tintinnids, by adhesion (Sugiyama and Anderson, 1997). In addition to predation, many spumellarian species dwelling in the sunlit ocean harbour photosynthetic algal symbionts, mainly identified as dinoflagellates (Probert et al., 2014; Yuasa et al., 2016; Zhang et al., 2018).

Radiolaria classifications are historically based on morphological criteria relying essentially on the initial spicular system; an early developed skeletal structure considered to be the foundation of the systematics at family and higher levels (De Wever et al., 2001). Traditionally, radiolarians with concentric structures and spherical or radial symmetry have been classified as Spumellaria or Entactinaria depending on the absence or presence, respectively, of such skeletal structure. Yet these taxonomic differences based on the initial spicular system have been recently questioned, finding molecular evidences for the polyphyly of Entactinaria among Rhizaria (Nakamura et al., 2020). The latest spumellarian classification attempted to merge the extensive morphological criteria (De Wever et al., 2001; O’Dogherty et al., 2009; Noble et al., 2017) with recent rDNA molecular studies (Matsuzaki et al., 2015). Outcome from this later work, the current classification scheme describes 9 extant superfamilies: Actinommoidea (Haeckel, 1862; O’Dogherty, 1994), Hexastyloidea (Haeckel, 1882), Liosphaeroidea (Haeckel, 1882), Lithelioidea (Haeckel, 1862; Petrushevskaya, 1975), Pylonioidea (Haeckel, 1882; Dumitrica, 1989), Saturnalioidea (Deflandre, 1953), Spongodiscoidea (Haeckel, 1862; De Wever et al., 2001), Sponguroidea (Haeckel, 1862; De Wever et al., 2001), Stylodictyoidea (Haeckel, 1882; Matsuzaki et al., 2015); and two undetermined families (Heliodiscidae, Haeckel, 1887; De Wever et al., 2001; and Spongosphaeridae Haeckel, 1862). In Matsuzaki et al. (2015) the authors stated the “overwhelmingly artificial” spumellarian classification outweighed by morphological criteria in regards to the little amount of molecular data available.

Few studies have explored the genetic diversity of Spumellaria, essentially unveiling relationships among higher rank taxonomical groups (Kunitomo et al., 2006; Yuasa et al., 2009; Krabberød et al., 2011; Ishitani et al., 2012). To date, with a total of 35 sequences from morphologically described specimens, covering 5 out of the 9 superfamilies described, mismatches between morphological and molecular characters are reported for two main groups of Spumellaria among which different innermost shell structure are found (Yuasa et al., 2009; Ishitani et al., 2012). Despite such important insights, the understanding of the inner structure relationships with the rest of the families remains elusive. Time-calibration of radiolarian phylogenies thanks to the fossil record allows a better understanding of relationships among extinct groups along with a contextualized evolutionary history (Decelle et al., 2012a; Lewitus et al., 2018; Sandin et al., 2019). In addition, acquisition of single cell reference DNA barcodes morphologically described establish the basis for further molecular ecology surveys, inferring the actual diversity and ecology in the nowadays oceans (Decelle et al., 2013; Nitsche et al., 2016; Biard et al., 2017).

Here we present an integrative morpho-molecular classification of Spumellaria obtained by combining ribosomal DNA taxonomical marker genes (18S and 28S partial rDNA) and imaging techniques (light and scanning electron microscopy). Phylogenetic placement of environmental sequences provided insights in the extant genetic diversity of Spumellaria in the contemporary oceans, allowing access to undescribed diversity. In addition, the extensive fossil record of Spumellaria allowed calibrating our phylogenetic analysis in time and inferring their evolutionary history contextualized with global scale geological and environmental changes from the Cambrian to present ecosystems.

## Material and Methods

### Sampling and single cell isolation

Plankton samples were collected: 1- off Sendai (11 samples: 38° 0’ 28.8’’ N, 142° 0’ 28.8’’ E), 2-Sesoko (8 samples: 26° 48’ 43.2’’ N, 73° 58’ 58.8’’ E), 3-the Southwest Islands, South of Japan (2 samples: 28° 14’ 49.2’’ N, 129° 5’ 27.6’’ E) 4-in the bay of Villefranche-sur-Mer (38 samples: 43° 40’ 51.6’’ N, 7° 19’ 40.8’’ E) and 5-in the western Mediterranean Sea (9 samples: MOOSE-GE cruise) by net tows (20, 64 or 200 μm), Vertical Multiple-opening Plankton Sampler (VMPS) or Bongo net (24-200 μm). Samples related metadata can be found in the RENKAN database (http://abims.sb-roscoff.fr/renkan). Spumellarian specimens were individually handpicked using Pasteur pipettes from the plankton tow sample and transferred 3 to 4 times into 0.2 μm filtered seawater to allow self-cleaning from debris, particles attached to the cell or prey digestion. Images of live specimens were taken under an inverted microscope and thereafter transfer into 1.5 ml Eppendorf tubes containing 50 μl of molecular grade absolute ethanol and stored at −20 °C until DNA extraction.

### DNA extraction, amplification and sequencing

DNA was extracted using the MasterPure Complete DNA and RNA Purification Kit (Epicentre) following manufacturer’s instructions. Both 18S rDNA and partial 28S rDNA genes (D1 and D2 regions) were amplified by Polymerase Chain Reaction (PCR) using Radiolaria and Spumellaria specific and general primers (Table 1). For further details about rDNA amplification see: dx.doi.org/10.17504/protocols.io.xwvfpe6. PCR amplicons were visualized on 1% agarose gel stained with ethidium bromide. Positive reactions were purified using the Nucleospin Gel and PCR Clean up kit (Macherey Nagel), following manufacturer’s instructions and sent to Macrogen Europe for sequencing.

**Table 1.**
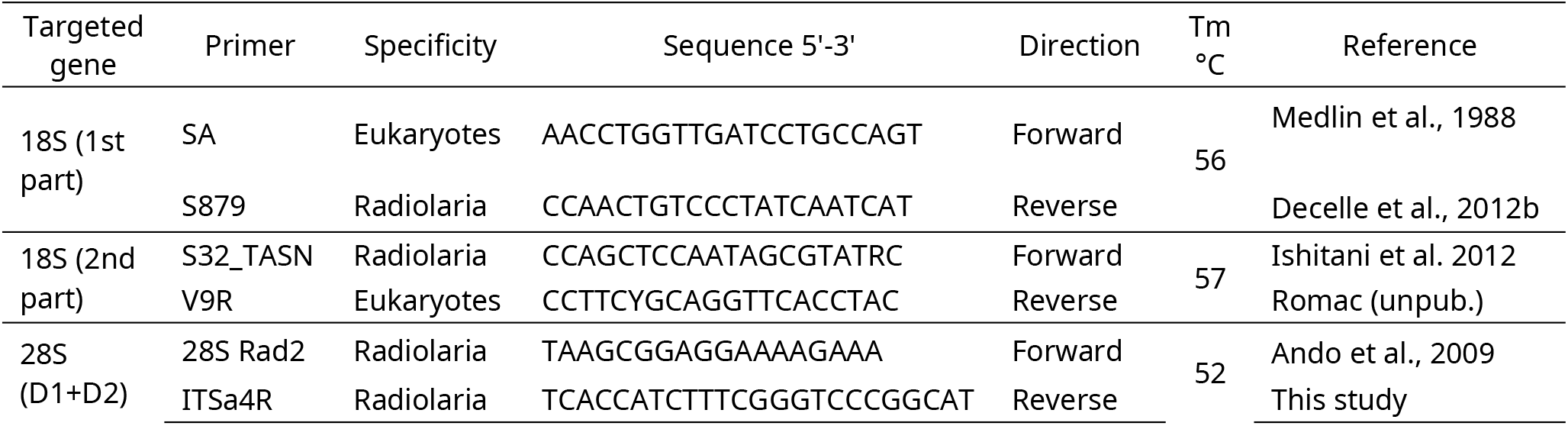
Primer sequences and temperatures used for DNA amplification and sequencing.

### Scanning Electron Microscopy (SEM)

After DNA extraction, spumellarian skeletons were recovered from the eluted pellet and handpicked under binoculars or inverted microscope. After cleaning and preparing the skeletons (detailed protocol in dx.doi.org/10.17504/protocols.io.ug9etz6) images were taken with a FEI Phenom tabletop Scanning Electron Microscope (FEI technologies).

### Single cell morphological identification

Spumellaria specimens were identified at the species level, referring to pictures of holotypes, through observation of live images and analysis of the skeleton by scanning electron microscopy when available (see Supplementary material Table S1 for taxonomic authority of specimens included in our study). Further details in taxonomic assignment of the specimens can be found in the Material and Methods of Sandin et al. (2019).

### Phylogenetic analyses

After sequencing, forward and reverse sequences were checked and assembled using ChromasPro software version 2.1.4 (2017). Sequences were compared to the GenBank reference database (GenBank) using the *BLAST search* tool integrated in ChromasPro to discriminate radiolarian sequences from possible contamination. Presence of chimeras, at a frequency as high as 13-25%, was detected by mothur v.1.39.3 (Schloss et al., 2009) against previously available reference sequences of Spumellaria. Sequences not detected as chimeras, were in turn included in our reference database for future phylogenetic reconstruction and chimeric analysis.

Two different datasets for each genetic marker (18S rDNA gene and partial 28S rDNA gene) were aligned separately using MAFFT v7.395 (Katoh and Standley, 2013) with a L-INS-i algorithm (‘--localpair’) and 1000 iterative refinement cycles for high accuracy. Each alignment was manually checked in SeaView version 4.6.1 (Gouy et al., 2010) and trimmed automatically using trimal (Capella-Gutiérrez et al., 2009) with a 30% gap threshold. For both genes, the 18S rDNA (133 taxa, 1789 positions) and the 28S rDNA (55 taxa, 746 positions), phylogenetic analyses were performed independently. The best nucleotide substitution model was chosen following the corrected Akaike Information Criterion (AICc) using the *modelTest* function implemented in the R version 3.6.0 (R Core Team, 2014) package *phangorn* version 2.5.5 (Schliep, 2011). The obtained model (General Time Reversible with Gama distribution and proportion of Invariable sites, GTR+G+I) was applied to each data set in R upon the packages *APE* version 5.3 (Paradis et al., 2004) and *phangorn* version 2.5.5 (Schliep, 2011). A Maximun Likelihood (ML) method (Felsenstein, 1981) with 10 000 replicates of bootstrap (Felsenstein, 1985) was performed to infer phylogenies.

Despite specific discrepancies in the topology of the two analysis (Supplementary material Fig. S1), the two genes were concatenated in order to increase taxonomic coverage and improve phylogenetic resolution. A final data set containing 133 taxa and 2535 positions was used to infer phylogenies following the previous methodology. Fourteen sequences of Nassellaria were assembled to form the outgroup as seen in previous classifications to be the sister clade of spumellaria (eg; Krabberød et al., 2011; Cavalier-Smith et al., 2018). The best model obtained was GTR+G+I, with 4 intervals of the discrete gamma distribution, and a ML method with 100 000 bootstraps were performed. In parallel, a Bayesian analysis was performed using BEAST version 1.8.4 (Drummond et al., 2012) with the same model parameters over 100 million generations sampled every 1000 states, to check the consistency of the topology and to calculate posterior probabilities. Final tree was visualized and edited with FigTree version 1.4.3 (Rambaut 2016). All sequences obtained in this study and used for the phylogeny were submitted in GenBank under the accession numbers: MT670437 - MT670549.

Morphological observations performed with light and scanning electron microscopy (Supplementary material Fig. S2) allowed assigning these sequences to 10 super-families (Actinommoidea, Hexastyloidea, Liosphaeroidea, Lithelioidea, Lithocyclioidea, Pylonioidea, Spongodiscoidea, Spongopyloidea, Spongosphaeroidea and Stylodictyoidea) and 2 super-families considered to belong to Entactinaria (Centrocuboidea and Rhizosphaeridae).

### Molecular clock analyses

Molecular clock estimates were performed according to Sandin et al. (2019). Nine nodes were chosen to carry out the calibration. The selection of these nodes is explained below from the oldest to the newest calibration age, and given the name of the node for the taxa they cover:

1. Root: The calibration for the root of the tree corresponds to the hypothesized last common ancestor between Nassellaria and Spumellaria (De Wever et al., 2001). A uniform distribution (U) with a minimum bound of 300 million years ago (Ma) and a maximum of 800 Ma (U[300, 800]) was set to allow uncertainty in the diversification of both groups and to establish a threshold restricting the range of solutions for the entire tree.
2. Nassellaria N(410, 20): The outgroup of the phylogeny is calibrated based in a consensus between the first appearance of nassellarian-like fossils in the Upper Devonian (ca. 372.2 Ma) in the fossil record (Cheng, 1986) and the first diversification of Nassellaria dated with previous analysis of the molecular clock (ca. 423 Ma; 95% HPD: 500-342 Ma; Sandin et al., 2019). Therefore, the node was normally distributed (N) with a mean of 410 and a standard deviation of 20: N(410, 20).
3. Spumellaria U[700, 200]: Recent studies have found the oldest spumellarian representatives in the early middle Cambrian (Zhang and Feng, 2019). Yet, De Wever et al. (2001) have argued that the initial spicular system may be subjected to preservation bias. Since many spumellarians and entactinarians (presence of the initial spicular system or not, respectively) are superficially similar and it is not possible to distinguish from one another (Suzuki and Oba, 2015), a uniform distribution was set to allow uncertainty in the diversification of Spumellaria.
4. Hexastyloidea N(242, 10): The family Hexastylidae is the first representative of the superfamily Hexastyloidea and has its first appearance in the fossil record in the Middle Triassic (Late Anisian: ca. 246.8 - 241.5 Ma; O’Dogherty et al., 2011).
5. Liosphaeroidea N(233, 20): The Liosphaeroidea seems to have appeared in the Triassic (Pessagno and Blome, 1980; De Wever et al., 2001). Although it is not sure whether there is a continuity between members from the Mesozoic and from the Cenozoic (Matsuzaki et al., 2015).
6. Actinommidae N(170, 20): The family Actinommidae appears for the first time in the fossil record in the Middle Jurassic (Aalenian: ca. 174.2-170.3 Ma; O’Dogherty et al., 2011) yet some morphologies from the Triassic resemblance this Superfamily (De Wever et al., 2001).
7. Rhizosphaeroidea N(148, 10): The Rhizosphaeridae appeared during the late Jurassic (Tithonian: ca. 145-152.1 Ma) in the fossil record (Petrushevskaya, 1975; De Wever et al., 2001; Afanasieva and Amon, 2006; Dumitrica, 2017).
8. Pylonioidea N(97, 10): The family Larnacillidae are the first representatives of the superfamily Pylonioidea appearing at the beginning of the Late Cretaceous (Cenomanian: ca. 100.5-93.9 Ma; De Wever et al., 2001, 2003; Afanasieva and Amon, 2006).
9. Coccodiscoidea N(45, 10): The family Coccodiscoidea appears for the first time in the fossil record in the Early Eocene (De Wever et al., 2001; Afanasieva and Amon, 2006).

### Post hoc analyses

Two different analyses were performed *a posteriori*: a diversification of taxa over time (Lineages Through Time: LTT) and an ancestral state reconstruction. The former analysis was carried out with the *ltt.plot* function implemented in the package *APE* (Paradis et al., 2004) upon the tree obtained by the molecular dating analyses after removing the outgroup. The second analysis used the resulting phylogenetic tree to infer the evolution of morphological characters. A numerical value was assigned to each state of a character trait, being 0 for the considered ancestral state, and 1 to 3 for the presumed divergence state. In total 6 traits were considered (Supplementary Material Table S2): the skeleton shape, the symmetry, the central structure, the internal cavity (outside of the central structure), the number of spines arisen from the central structure and the number of distinctive cortical shells. Once the character matrix was established a parsimony ancestral state reconstruction was performed to every character independently in Mesquite version 3.2 (Madison & Madison, 2017).

### Environmental sequences

Each of the reference 18S rDNA and partial 28S rDNA spumellarian sequence available in our study was compared with publicly available environmental sequences in GenBank (NCBI) using BLAST (as of May 2019). It allowed estimating the environmental genetic diversity of Spumellaria and to assess the genetic coverage of our phylogenetic tree. A total of 1171 and 6 environmental sequences affiliated to Spumellaria were retrieved for the genes 18S rDNA and 28S rDNA respectively. Out of the six 28S rDNA sequences, 4 were covering the full rDNA gene and also found in the 18S rDNA survey. These 4 sequences along with the 8 sequences already included in the phylogeny (2 in clade H and 8 in clade I) were therefore removed to avoid duplicates. Final dataset containing 1165 environmental sequences were aligned against the reference alignment used for the phylogenetic analysis using MAFFT v7.395 (Katoh and Standley, 2013) with the “--add” option and placed in our reference phylogenetic tree using the pplacer software (Matsen et al., 2010). A RAxML (GTR+G+I) tree was built for the placement of the sequences with a rapid bootstrap analysis and search for best-scoring ML tree and 1000 bootstraps.

In order to discriminate pseudo-genes among environmental clades due to a high frequency of chimeras encountered and long phylogenetic branches, two preliminary analysis were performed: a Shannon entropy analysis and an estimation of the GC content. Shannon entropy was calculated for every position of the resulting alignment matrix, without considering gaps due to differences in length of environmental sequences (script available on https://github.com/MiguelMSandin/DNA-alignment-entropy). The GC content was estimated for every sequence independently as follows: (G+C)/(G+C+A+T), where G, C, A and T are the number of guanines, cytosines, adenines and thymines respectively. Analysis were performed in R version 3.6.0 (R Core Team, 2014) and plotted with the package *ggplot2* version 3.2.1 (Wickham, 2016).

## Results

### Comparative molecular phylogeny and morphological taxonomy

Our final molecular phylogeny is composed of 133 distinct spumellarian sequences of the full 18S rDNA gene, completed with 55 sequences of the partial 28S (D1 & D2 regions) rDNA gene (Supplementary material Table S1). From the 133 final sequences of the 18S rDNA, 67 were obtained in this study, 58 were previously available sequences associated to a morphologically described specimen and 8 were environmental sequences not affiliated to any morphology, added to increase phylogenetic support in poorly represented clades. Regarding the partial 28S rDNA, a total of 37 sequences were obtained in this study and 18 were previously available. The final alignment matrix has 25.45% of invariant sites.

Phylogenetic analyses show 13 different clades (Clades A, B, C, D, E, F, G, H, I, J, K, L and M) clearly differentiated by BS and PP values distributed in 4 main lineages (I, II, III, and IV) showing high BS (>97) and PP (1) and consistent position across the different phylogenetic analyses (Fig. 1). In general, morphological classification agrees with molecular phylogeny at the clade level. Considered together, morphological similarities and molecular phylogenetic support established super-families Hexastyloidea (spread in clades A, B and C), Spongosphaeroidea (Clade D), Lithocyclioidea (Clade E1) and Spongodiscoidea (Clade E2 and E3) in lineage I; Liosphaeroidea (Clade F) in lineage II; Rhizosphaeroidea (Clade G) and Centrocuboidea (Clades H and I) in Lineage III and Actinommoidea (Clade K), Stylodictyoidea (Clade J), Lithelioidea (Clade L1), Spongopyloidea (Clade L2) and Pylonioidea (Clade M) in lineage IV. All these morpho-molecular clades are highly supported in both the 18S rDNA and 28S rDNA gene phylogenies, but clades B and E in the 28S rDNA gene phylogeny (Supplementary material Figure S1). In contrast to the clade definition, the general topology and relationships between clades slightly disagree between the 18S rDNA and the 28S rDNA gene phylogenies. Such discrepancies are basically due to the variable position of Clade F that in the 18S rDNA gene phylogeny appears basal to all clades and in the 28S rDNA gene appears basal to clades J, K, L and M, weakly supported as a sister group of clades H and G. Another example is, clades B, C, D and E, which are in the same highly supported group in the 18S rDNA gene phylogeny, whereas in the 28S rDNA gene phylogeny these clades appear scattered at basal positions regarding the rest of the clades. In the concatenation of the two genes clade F appears highly supported as a group with the clades G to M (Fig. 1).

**Figure 1.**
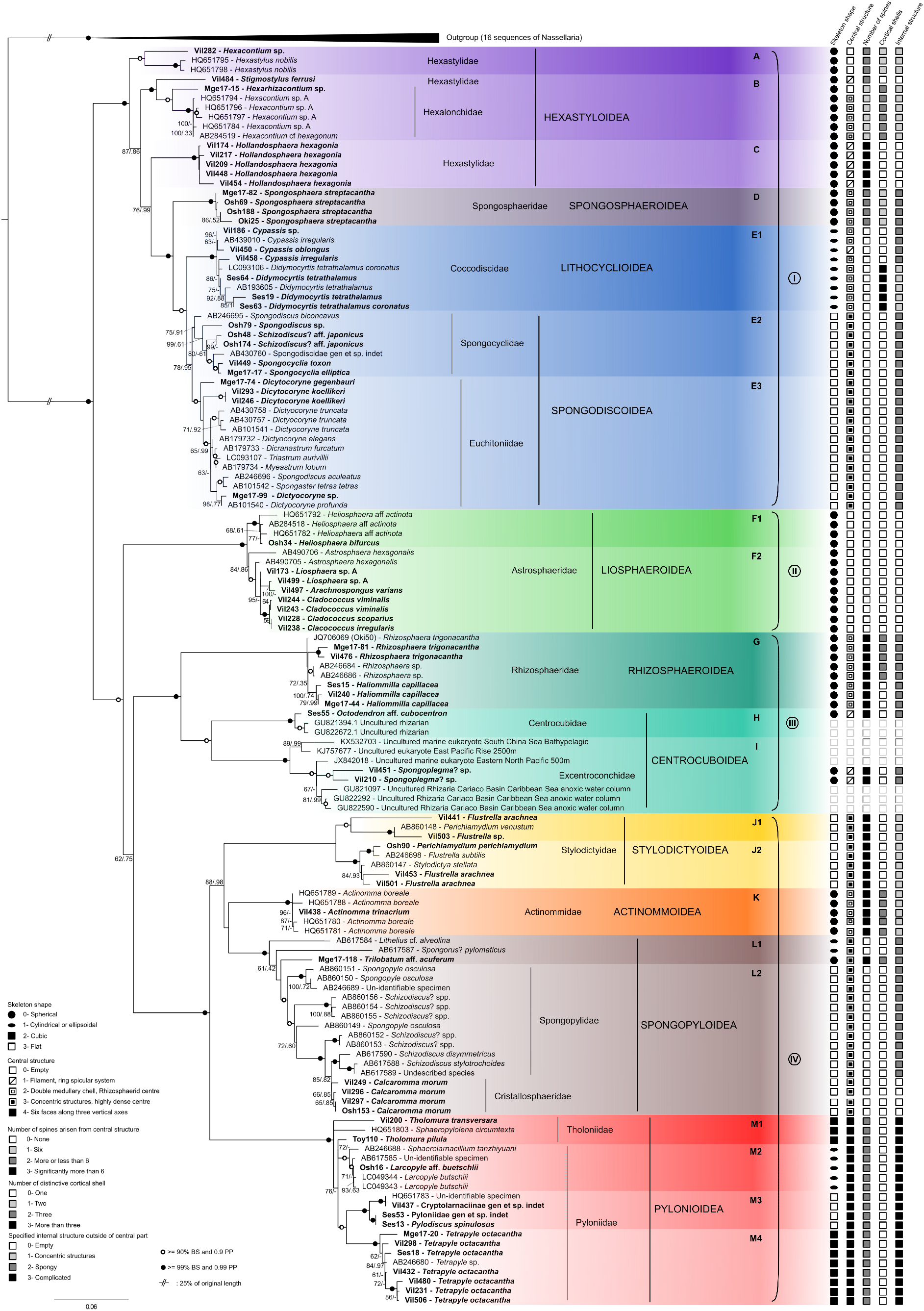
Molecular phylogeny of Spumellaria inferred from the concatenated complete 18S and partial 28S (D1-D2 regions) rDNA genes (135 taxa and 2459 aligned positions). The tree was obtained by using a phylogenetic Maximum likelihood method implemented using the GTR + γ + I model of sequence evolution. PhyML bootstrap values (100 000 replicates, BS) and posterior probabilities (PP) are shown at the nodes (BS/PP). Black circles indicate BS ≥ 99% and PP ≥ 0.99. Hollow circles indicate BS ≥ 90% and PP ≥ 0.90. Sequences obtained in this study are shown in bold. Thirteen main clades are defined based on statistical support and morphological criteria (A, B, C, D, E, F, G, H, I, J, K, L, M), divided in 4 main lineages (I, II, III, IV). For each morpho-molecular clade, capital letters represent the Superfamily name and small letters the Family name. Main skeletal traits are identified on the right for each spumellarian specimen. Fourteen Nassellaria sequences were assembled as outgroup. Branches with a double barred symbol are fourfold reduced for clarity.

Lineage I includes clades A, B, C, D and E and is phylogenetically distant to the other 3 lineages. Lineage I is characterized by the presence of one or two concentric shells or a full spongy test, sometimes non-distinguishable, where the main spines grow from the innermost shell and go through the outermost shell, when present (Fig. 2. A-E). Only clade A and B shows two concentric shells with six primary spines from the center (Fig. 2. A-B). Clade A is composed of 1 novel and 2 previously available sequences and a common characteristic is the fragile and hexagonal mesh constituting the outermost shell (Fig. 2.A). Clade B is composed of 2 novel and 5 previously available sequences clustered with high BS and PP values despite their phylogenetic distance (Fig. 1). All specimens of clade B show a more irregular and thicker mesh of the shell compare to that of clade A, and, when present, very thin spines (generally called as by-spines) coming out from all the surface of the shell (Fig. 2. Ba and Bb). Clade C is composed of 5 novel sequences showing a large single double layered shell with no main spines but a large abundance of long and thin spines (Fig. 2. C). These three clades agree with the definition of the Superfamily Hexastyloidea (Haeckel, 1882; Matsuzaki et al., 2015) displaying an innermost shell that, if present, always shows pyriform with six radial beams. Three different families within Hexastyloidea correspond to: Hexastylidae (Haeckel, 1887) in clade A, Hexalonchidae (Haeckel, 1882) in clade B and Hollandosphaeridae (Deflandre, 1973) in clade C. The innermost shell if present always shows pyriform with six radial beams. Clades D and E are the more distal clades in this lineage and highly supported together (100 BS and 1 PP). The presence of spongy structures differentiates these two clades from the others of lineage I (Fig. 2. D-E). Clade D is composed of 4 novel sequences obtained in this study. All specimens of this clade show a single, very small spherical shell where several long and three bladed spines grow interconnected by a spongy mesh (Fig. 2. D). This definition agrees with the genus *Spongosphaera* belonging to the Superfamily Spongosphaeroidea (Haeckel, 1862). The last clade of Lineage I (Clade E) is composed of 30 sequences of which 15 are novel. Members of this clade tend to lose the concentric symmetry towards a cylindrical to ellipsoidal (sub-clade E1; Fig. 2. E1a and E1b) or flat symmetry (sub-clade E2, Fig. 2. E2a and Eb; and Sub-clade E3, Fig. 2. E3), and so does the spines and the shell. Spongy structures take more importance complicating the inner structure. These morphologies agree with the definition of the Family Artiscinae (Haeckel, 1882) for E1 and the Superfamily Spongodiscoidea (Haeckel, 1862; De Wever et al., 2001) for E2 and E3. Within the latest clades there are representatives of two families Spongobrachiidae (Haeckel, 1882) with moderate BS and PP values in sub-clade E2 and Eucthitoniidae (Haeckel, 1882) highly supported in sub-clade E3.

**Figure 2.**
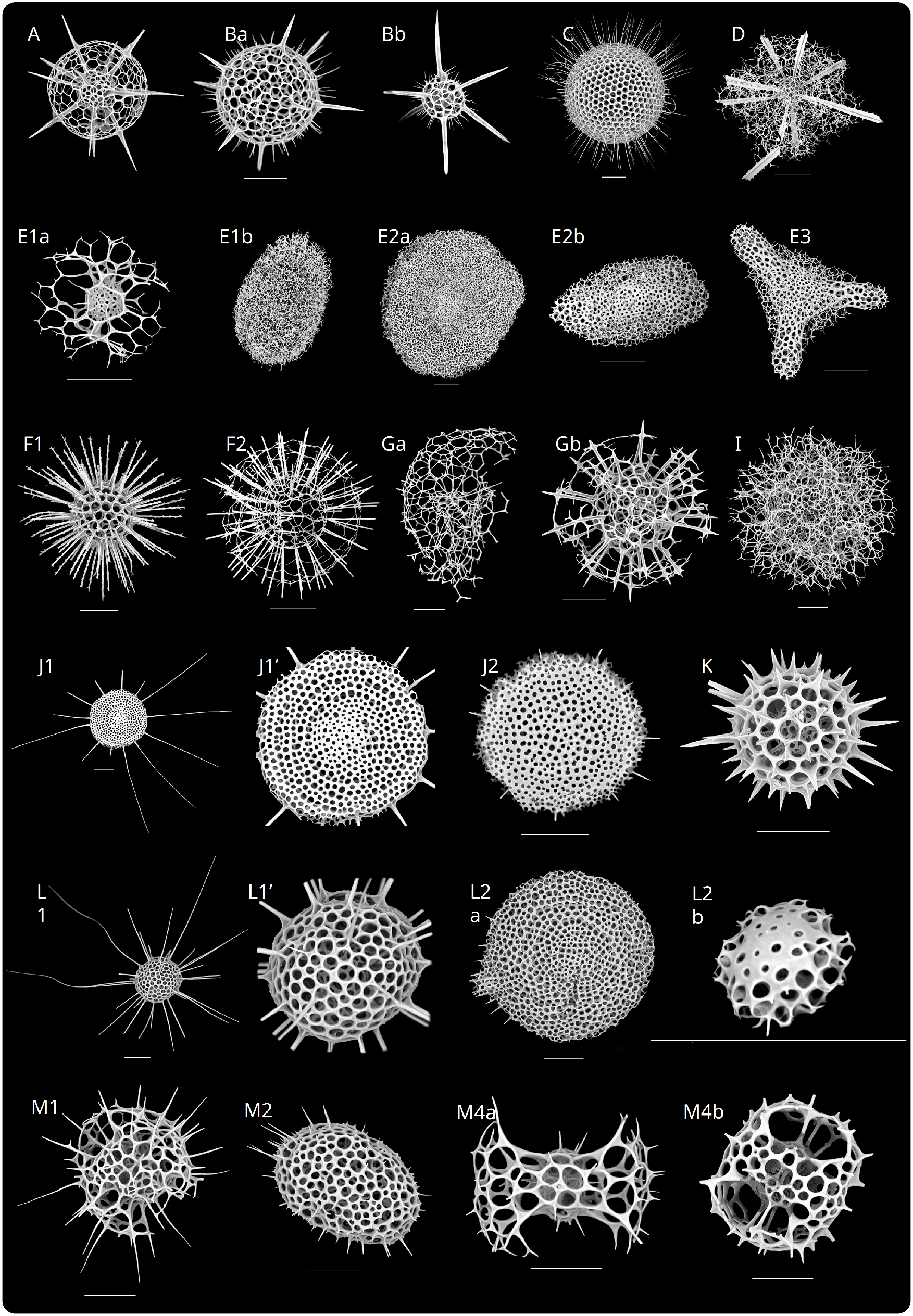
Scanning Electron Microscopy (SEM) images of Spumellaria specimens used in this study for phylogenetic analysis or morphologically related to one of the morpho-molecular clades of Fig. 1. Letters correspond to its phylogenetic clade in Fig. 1. Scale bars = 50μm. (**A**) Osh194: *Hexacontium sp.* (specimen not included in phylogeny). (**Ba**) Vil484: *Stigmostylus ferrusi*. (**Bb**) Mge17-15: *Hexacontella*? sp. (**C**) Vil174: *Hollandosphaera hexagonia*. (**D**) Mge17-82: *Spongosphaera streptacantha*. (**E1a**) Vil186: *Didymocyrtis* sp. (**E1b**) Vil458: *Cypassis irregularis*. (**E2a**) Osh48: Spongodiscidae gen et sp. Indet. A (**E2b**) Mge17-17: *Spongobracium ellipticum*. (**E3**) Vil246: *Dictyocoryne koellikeri*. (**F1**) Osh34: *Heliosphaera bifurcus*. (**F2**) Vil499: *Cladococcus* sp. A. (**Ga**) Vil240: *Haliommilla capillacea* (broken specimen). (**Gb**) Mge17-81: *Rhizosphaera trigonacantha*. (**I**) Vil210: *Plegmosphaeromma*? sp. (**J1**) Vil441: *Stylodictya arachnea*. (**J1’**) detail of Vil441. (**J2**) Osh90: *Stylodictya stellata*. (**K**) Vil438: *Actinomma trinacrium*. (**L1**) Mge17-118: *Trilobatum* aff. *acuferum*. (**L1’**) detail of Mge17-118. (**L2a**) Osh191: *Schizodiscus* spp. (specimen not included in phylogeny). (**L2b**) Vil296: *Calcaromma morum*. (**M1**) Vil200: *Tholomura transversara*. (**M2**) Osh16: *Larcopyle* aff. *buetschlii*. (**M4a**) Mge17-20: *Tetrapyle octacantha*. (**M4b**) Vil452: *Tetrapyle octacantha (specimen not included in phylogeny)*.

Lineage II, including clade F alone, is composed of 13 sequences of which 5 were previously available and 8 are novel. All specimens within this clade are characterized by one large and hollow shell where long spines grow from its surface (Fig. 2. F1 and F2), that agree with the definition of the Superfamily Liosphaeroidea (Haeckel, 1882). Members of the sub-clade F1 (Fig. 2. F1) share a robust shell compared to that of sub-clade F2 (Fig. 2. F2) and short spicules are coming out of the spines. Those specimens agree with the definition of the family Astrosphaeridae (Haeckel, 1887; Matsuzaki et al., 2015) in the large hollow and fragile shell with the presence of long or short spicules coming out at the far end of the spines.

Within Lineage III, including clades G, H and I, Clade G is represented by 5 novel and 3 previously available sequences. Members of this clade have a very unique innermost shell, named “rhizosphaerid-type microsphere” or “microbursa” (Dumitrica et al. 2010) with a large and fragile spherical skeleton (Fig. 2. Ga) or several cortical shells with a robust skeleton (Fig. 2. Gb). Either morphology agree with the definition of the family Rhizosphaeridae (Hollande and Enjumet, 1960; Dumitrica, 2017). Clade H is composed by 1 novel sequence with a morphological reference (Ses55, Supplementary material figure S2) and 2 environmental sequences not morphologically described. The picture available for this specimen lacks taxonomic resolution, although the skeletal structure resemble that shown in Aita et al. (2009, pl. 23, fig. 3) specified as *Centrocubus cladostylus* (Haeckel, 1887). This species has characteristic interconnected spines and the highly supported position between clade G and I agrees with the definition of the family Centrocubidae (Hollande and Enjumet, 1960; emend. Dumitrica, 1994). The last clade of this lineage (Clade I) is represented by 2 novel sequences obtained in this study and 6 environmental sequences. Specimens of this clade show only spongy structures with radiated fibers and there is no evidence of either shell or main spines (Fig. 2. I).

**Figure 3.**
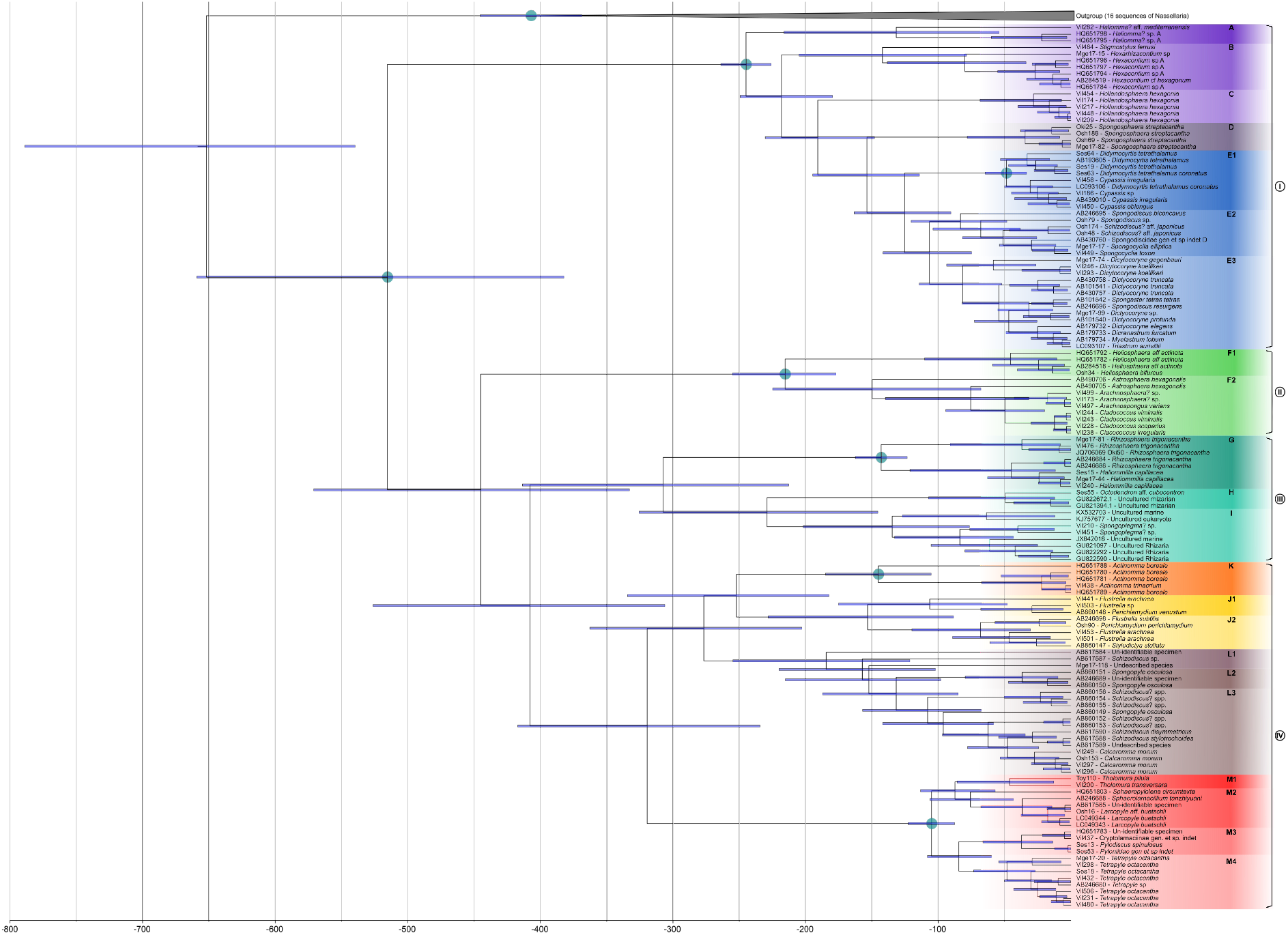
Time-calibrated tree (Molecular clock) of Spumellaria, based on alignment matrix used for phylogenetic analyses. Node divergences were estimated with a Bayesian relaxed clock model and the GTR + γ + I evolutionary model, implemented in the software package BEAST. Nine different nodes were selected for the calibration (blue dots). Blue bars indicate the 95% highest posterior density (HPD) intervals of the posterior probability distribution of node ages.

In Lineage IV, Clade J, K and L cluster together with relatively high BS (88) and PP (0.98) values as a sister group of clade M. Although relationships between clades J, K and L remain still elusive due to the short phylogenetic distance and the different topology when using a Maximum Likelihood (Fig. 1) or a Bayesian (Fig. 3) approach (BS: 49 between clade K and L and PP: 0.38 between clade J and K). All members of this Lineage share a very small spherical inner most shell with four or more radial beams connecting to the second shell, where 12 or more radial beams come out (Fig. 2. J-M). Lineage IV differs from Lineage I in the absence of discrete radial beams from the second or later shells. Clade J is characterized by 5 sequences obtained in this study and 3 previously available sequences. Members of this clade share a flattened skeleton with 2 or more concentric shells and radial spines from the innermost or second innermost shell (Fig. 2. J1, J1’ and J2), agreeing with the definition of the Superfamily Stylodictyoidea (Haeckel, 1882; Matsuzaki et al., 2015). Morphological differences between sub-clade J1 (Fig. 2. J1) and J2 (Fig. 2. J2) were not possible to determine, yet they have been separated into two different sub-clades due to high BS (96 in J1 and 100 in J2) and PP (0.98 in J1 and 1 in J2) values. Clade K contains 1 novel and 4 previously available sequences. All specimens of this clade have three spherical shells with more than eight main spines that extend from the inner shell (Fig. 2. K), representing the Superfamily Actinommoidea (Haeckel, 1862). Clade L is represented by 5 novel and 14 previously available sequences. Members of this clade are morphologically distant. Sub-clade L1 has a tightly concentric or coiled center with one thick protoplasmic pseudopodium (axopodium) (AB61787) or a rhomboidally inflated center (Fig. 2. L1 and L1’), which is not observed in any other Spumellaria. In the first case, this morphology is attributed to the Superfamily Sponguroidea, and in the second to Lithelioidea. Sub-clade L2 is represented by either: a flatten circular test with a tunnel-like pylome and a test comprised by a very densely concentric structure (Fig. 2. L2a) attributed to Spongodiscoidea; or by a spherical translucent protoplasm with an encrypted flat consolidated skeletal shell in distal position and star-like solid soluble materials (Fig. 2. L2b and Supplementary material Fig. S2) assigned to Collodaria. In Clade M there are 13 novel sequences and 7 previously available sequences. A morphological characteristic of this clade is the broken, or fenestrated, outermost shell with the presence of pyloniid central structure (Fig. 2. M), corresponding to the Superfamily Pylonioidea (Haeckel, 1882), thereafter, the symmetry and the different opening of the outermost shell distinguish the sub-clades. Yet, phylogenetic relationships within this clade remain elusive. The first sub-clade (M1) consists of three basal sequences poorly supported as a clade yet sharing a cubic morphology and the two opposite and closed gates (Fig. 2. M1), which agrees with the family Tholoniidae (Haeckel, 1887). The three other sub-clades have different morphologies (Fig. 2. M2; Ses13, Ses53 and Vil437, Supplementary material Fig. S2) agreeing with the definition of different families within Pylonioidea. Sub-clade M2 (Fig. 2. M2) has a spherical to ellipsoidal skeleton without distinctive openings. On the other side, sub-clades M3 and M4 are highly supported as a group with a flatten box-shaped skeleton for the former sub-clade (Ses13, Ses53 and Vil437, Supplementary material Fig. S2) and a cubic skeleton characterized by big openings of the second shell for the later sub-clade (Fig. 2. M4a and 4b).

### Molecular dating

The molecular clock dated the diversification between Nassellaria and Spumellaria (the root of the tree) at a median value of 651 Ma (95% Highest Posterior Density -HPD-: between 789 and 540 Ma) (Fig. 3). From here on, all dates are expressed as median values followed by the 95% HPD interval. The first diversification of Spumellaria happened around 515 (659-382) Ma, followed by a rapid branching of the lineages. Despite their dubious phylogenetic relationships, Lineage II splits apart 445 (571-332) Ma and Lineage III and IV diversified from each other 407 (526-306) Ma. The next diversification events correspond to the first radiation of Lineage IV 319 (417-234) Ma and that of Lineage III 307 (414-213) Ma. Within Lineage IV, the phylogenetic relationships are doubtful, yet two radiation events at 277 (363-203) Ma and 252 (334-182) Ma separate Clades J, K and L. Clade L is the first of these clades diversifying at 184 (255-121) Ma, followed by clade J 153 (228-88) Ma, clade K at 145 (185-105) Ma and Clade M at 105 (123-88) being the youngest clade of Lineage IV. Lineage III diverged soon after Lineage IV, yet the next ramification was between Clade G and Clade I 229 (326-145) Ma. It was not until 143 (162-123) and 135 (202-76) Ma when Clade G and I respectively radiate, and the latest diversification within this Lineage corresponds to Clade H 49 (107-12) Ma. Lineage I appeared 245 (264-226) Ma, and thereafter is characterized by a series of relatively continuous diversification events until the radiation of Clades B, A, E, D and C at 142 (205-79), 131 (216-54), 153 (163-90), 34 (78-8) and 28 (68-7) Ma respectively. Regarding Lineage II, it is the youngest of all the lineages appearing at 215 (255-177) Ma despite its early branching from any other lineage.

### Post-hoc analyses

The lineages through time analysis (Fig. 4) showed a classic exponential diversification slope, with a 0.0102 rate of speciation (Ln(lineages)·million years^−1^). The first diversification of extant Spumellaria corresponds to an early and fast divergence of the different lineages from ~515 to ~407 Ma. After that, the diversification of living groups remained standstill until ~313 Ma when Lineage IV and III radiated. From ~276 to ~215 Ma all different groups diversified: Clades K, J and L (within Lineage IV) split apart; Lineage I appeared and started a continued diversification; within Lineage III the three clades split apart; and Lineage II emerged. Another important diversification event happened from ~157 to ~125 Ma when Lineage IV greatly diversified, followed by Lineage I and II. During the following years there is a tiered diversification, where Clade M appeared in isolated events as well as the ramification of Clade L or the branching between sub-clades E2 and E3. Finally, from ~54 Ma on wards there was a continuous diversification where the rest of the clades appeared or keep diversifying. During this time lineage III diversified notably and so did lineage II.

**Figure 4.**
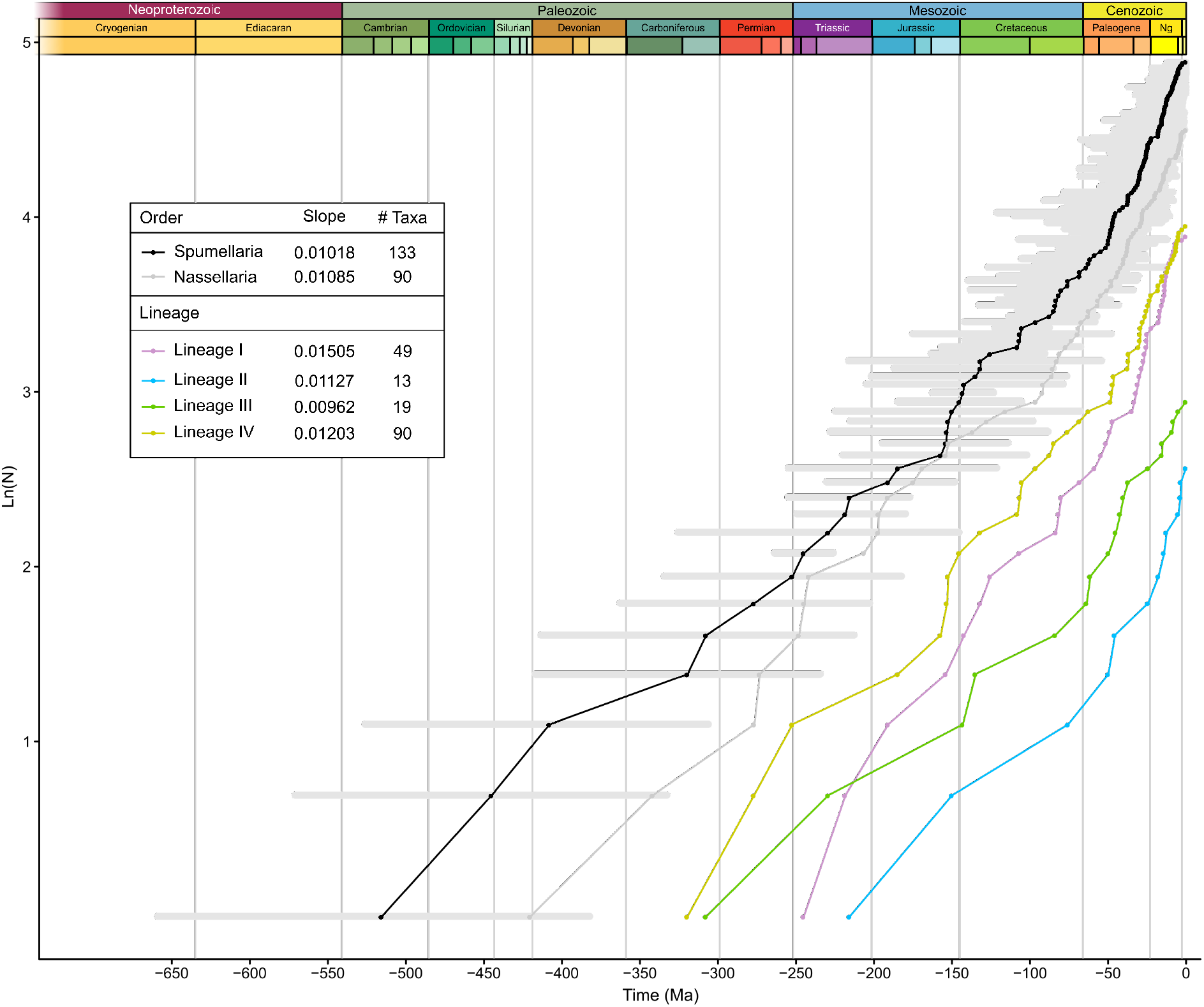
Lineages Through Time (LTT) analysis based on the molecular clock results for Spumellaria, removing the outgroup (in black) and of each lineage independently: lineage I (purple), lineage II (light green), lineage III (dark green) and lineage IV (orange) and of Nassellaria (data taken from Sandín et al., 2019). The y-axis represents the number of lineages (N) expressed in logarithmic (base e) scale (Ln(N)) and in the x-axis it is represented the time in million of years ago (Ma). Horizontal grey bars in black slope represent the 95% Highest Posterior Density (HPD) of molecular clock estimates.

The ancestral state reconstruction analysis (Supplementary Material Fig. S3) established a spherical skeleton shape with a line of symmetry for the common ancestor to all Spumellaria, with a central structure that can be either empty (state 0) or with an entactinarian-type microsphere or fused fibers or soft spongy system (state 1), but with a wide space in between the central structure and the cortical shell, without spines arising from it and one distinctive cortical shell. The common ancestor of Lineage I and that of Lineages II, III and IV share a spherical skeleton shape with a line of symmetry, a wide internal cavity and one cortical shell. The difference between these two ancestors is that the ancestor of lineage I is characterized by an entactinarian-type central structure or fused fibers or soft spongy system with more or less 6 spines, whereas that of lineages II, III, IV could also have an empty central structure and have no spines arising from the central structure. The common ancestor for Lineage II has the most ancestral state in every reconstructed trait (state 0). The common ancestor of lineages III and IV have also a spherical skeleton shape with a line of symmetry, like the common ancestor of these two lineages independently. Although the common ancestor of Lineages III and IV differs from that of Lineage II in a higher complexity of the central structure, the appearance of structures in the internal cavity, the presence of significantly more than 6 spines and that it could have 3 cortical shells. Lastly, the common ancestor of the lineage III and that of lineage IV could also have 1 or 3 cortical shells with significantly more than 6 radial spines. Yet in Lineage III the internal cavity gets more dense than that of Lineage IV, whereas the central structure tend to be entactinarian-type, fused fibers or soft spongy in Lineage III and that of the common ancestor of Lineage IV is undefined since it has equal probabilities every character state.

### Environmental genetic diversity of Spumellaria

A total of 1165 environmental sequences affiliated to Spumellaria, and retrieved from NCBI public database, were placed in our reference phylogenetic tree (Fig. 5; Supplementary material Table S3). From these, 617 were closely related to Clade F. Two other clades with a high number of environmental sequences related to are Clade I and Clade M, with 145 and 47 sequences respectively. While up to 40 sequences were scattered between clades E, L, B, G, D and J (17, 14, 3, 3, 1 and 1 respectively), no environmental sequences were clustered within clades A, C, H and K. The rest of the sequences (317) where distributed across the tree, mainly at basal nodes, with no possible assignation: 206 basal to clade F in lineage II, 70 in Lineage I (of which 41 basal to the lineage, 11 basal to clades C, D and E, 12 basal to clades D and E, 6 basal to clade B and 1 basal to clade C), 26 in lineage IV (of which 16 basal to clade J, 5 basal to clade M and 5 basal to the lineage), 4 in lineage III basal to clade G, 2 basal to lineages II, III and IV and 8 final sequences basal to all Spumellaria.

**Figure 5.**
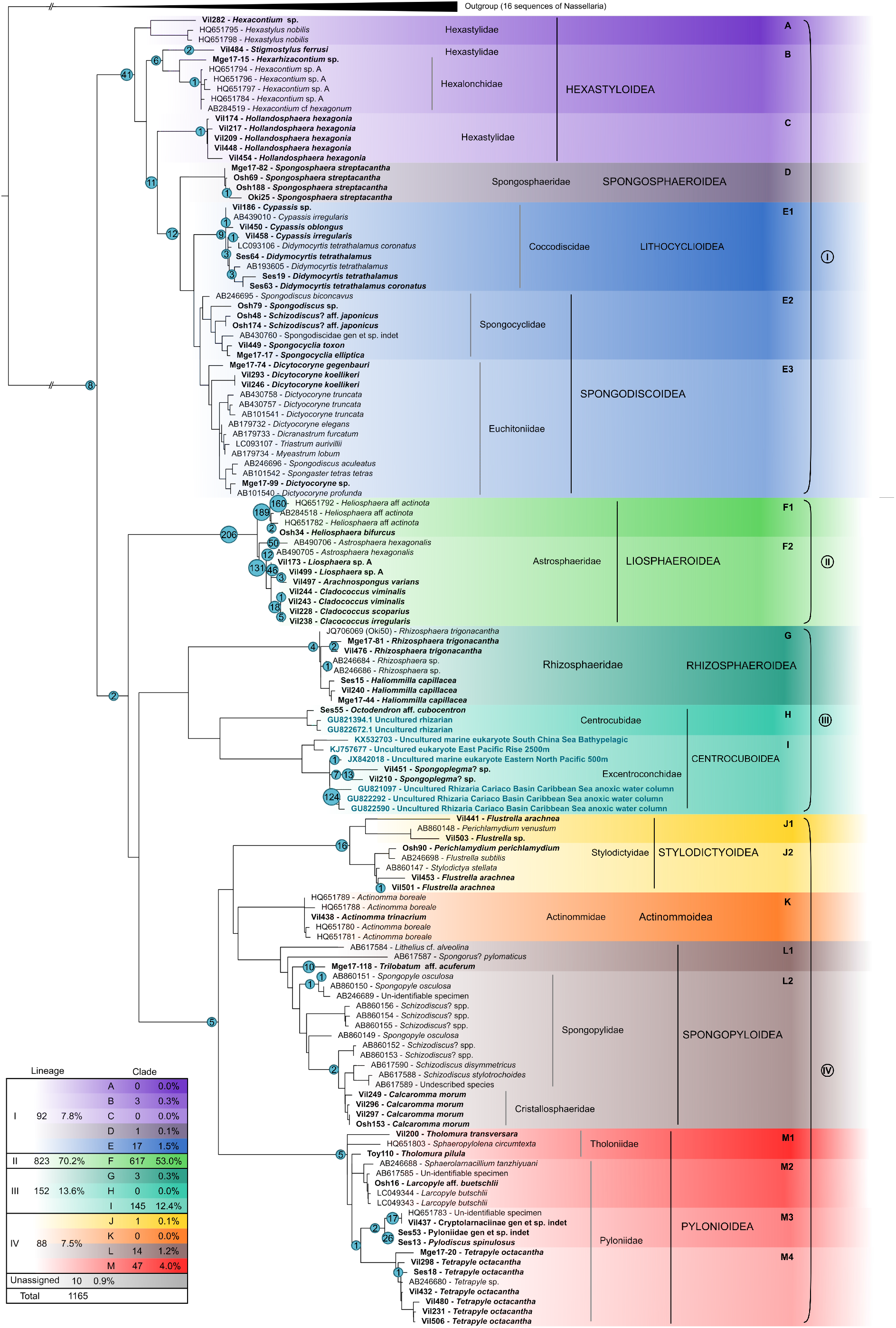
Pplacer phylogenetic placement of 1165 environmental sequences into a concatenated phylogenetic tree of Spumellaria (complete 18S + partial 28S rDNA genes). Numbers at nodes represent the amount of environmental sequences assigned to a branch or a node.

To go beyond rapid phylogenetic placement using the pplacer tool, these 317 environmental sequences dubiously assigned, were later included in a phylogenetic tree of the 18S rDNA gene (RAxML v8.2.10, GTR+G+I, with 1000 rapid bootstraps, Stamatakis, 2014) after removing chimeras (52 sequences were detected as chimeric) and suspicious sequences (11 sequences with multiple phylogenetic placements and/or long branches). Up to 6 different new clades were found (Env1 – 6; Fig. 6), 6 sequences scattered over the phylogenetic tree (1 related to Env2, 3 sequences related to Clade F and Env5 and 2 sequences closely related to Clade G) and 2 last sequences were clustered with high support basal to Clade M (Fig. 6; further details of sequence assignations can be found in Supplementary material Table S3). From these clades, Env5 gathers 149 sequences related to clade F in lineage II, yet low bootstrap values (51) avoid a formal assignation. Two clades (Env1 with 63 sequences and Env2 with 7) are found at basal positions of lineage I. Another environmental clade appears in lineage IV (Env4 with 16 sequences), and two final clades are found basal to lineages I, III and IV (Env6 with 8 sequences) and basal to lineage IV (Env3 with 3 sequences).

**Figure 6.**
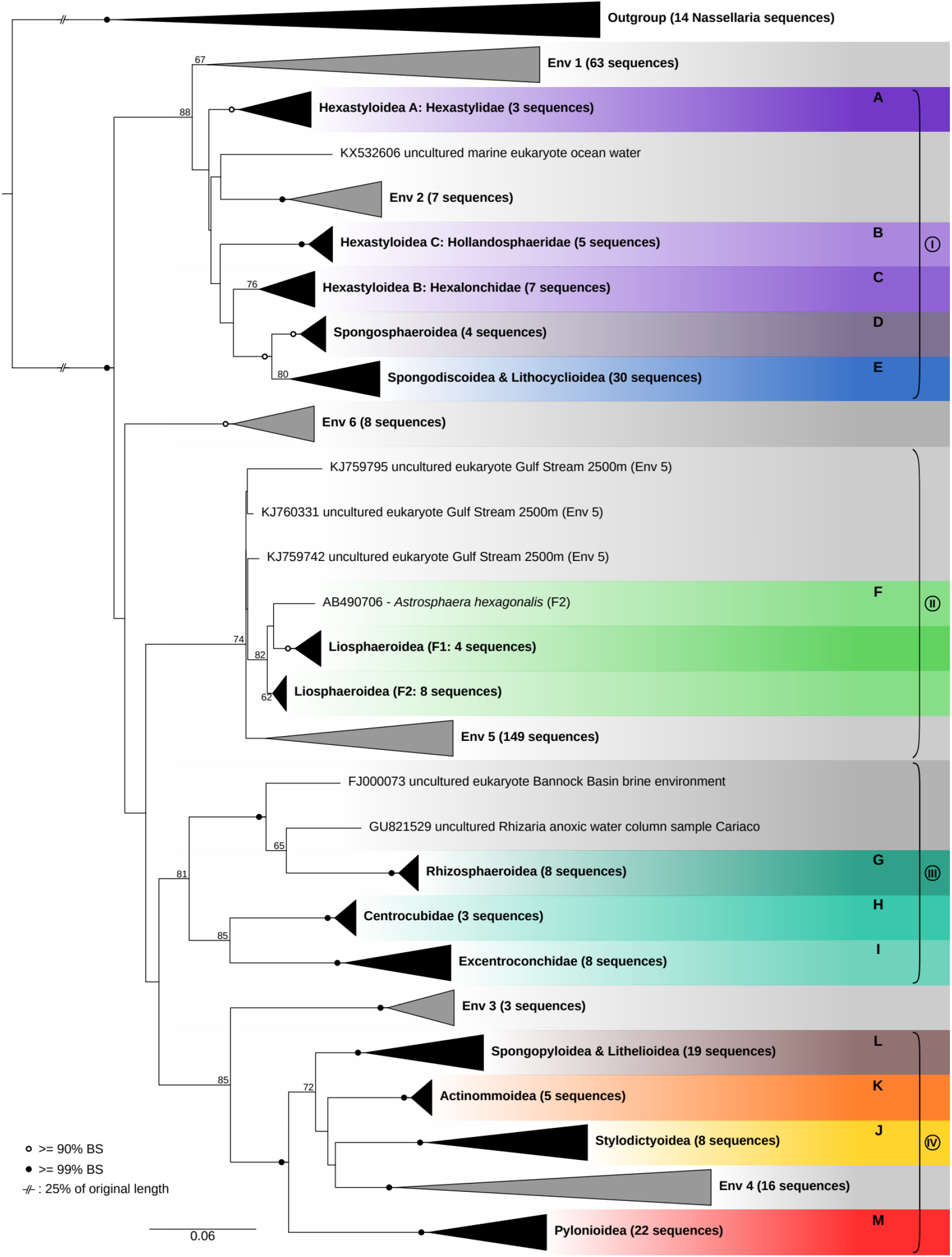
Molecular phylogeny of environmental sequences associated to Spumellaria. The tree was obtained by using 1000 rapid bootstrap RAxML GTR + γ model of sequence evolution. Black circles at nodes indicate BS ≥ 99% and hollow circles indicates BS ≥ 90%. BS lower than 60 were removed for clarity. Branches with a double barred symbol are fourfold reduced for clarity. Black triangles represent clades which morphology is known (Fig. 1, Fig. 2). Grey triangles represent environmental clades. Sequence composition of each environmental clade is described in Table S3.

In order to improve the understanding of the diversity of the environmental clades (Fig. 6) and try to explain the high chimeric ratio (~13-25%) observed, we performed a second phylogenetic analysis (RAxML v8.2.10, GTR+G+I, with 1000 rapid bootstraps, Stamatakis, 2014) including the dataset used to build fig. 6 and adding 9 sequences obtained from single-cell isolates but previously removed due to the contrasting morphological assignation in the clustered phylogenetic clade. It turns out that these 9 sequences clustered with moderate to high support among one of the environmental clades, clade Env1 (Fig. 7; Supplementary material Table S1). These 9 specimens show a morphology that matches clades E (Osh87 and Osh174), F (Osh156), G (Osh82, Osh97 and Osh114), I (Osh187), L (Osh116) and a last specimen with a morphology not covered in our morpho-molecular framework (Osh108). Our morpho-molecular classification show that each molecular clade has a clear and endorsed (2 or more specimens) morphological patterns distinct from one another. The combination of different morphologies within the same clade Env1 contrasts with this global pattern. In addition, one of the specimens from clade Env1 (Osh174) is also found within clade E. Although in clade E it is composed by the concatenation of the first part of the 18S rDNA and the 28S rDNA gene (1489 bp in total) and that of clade Env1 by the second part of the 18S rDNA gene (871 bp).

**Figure 7.**
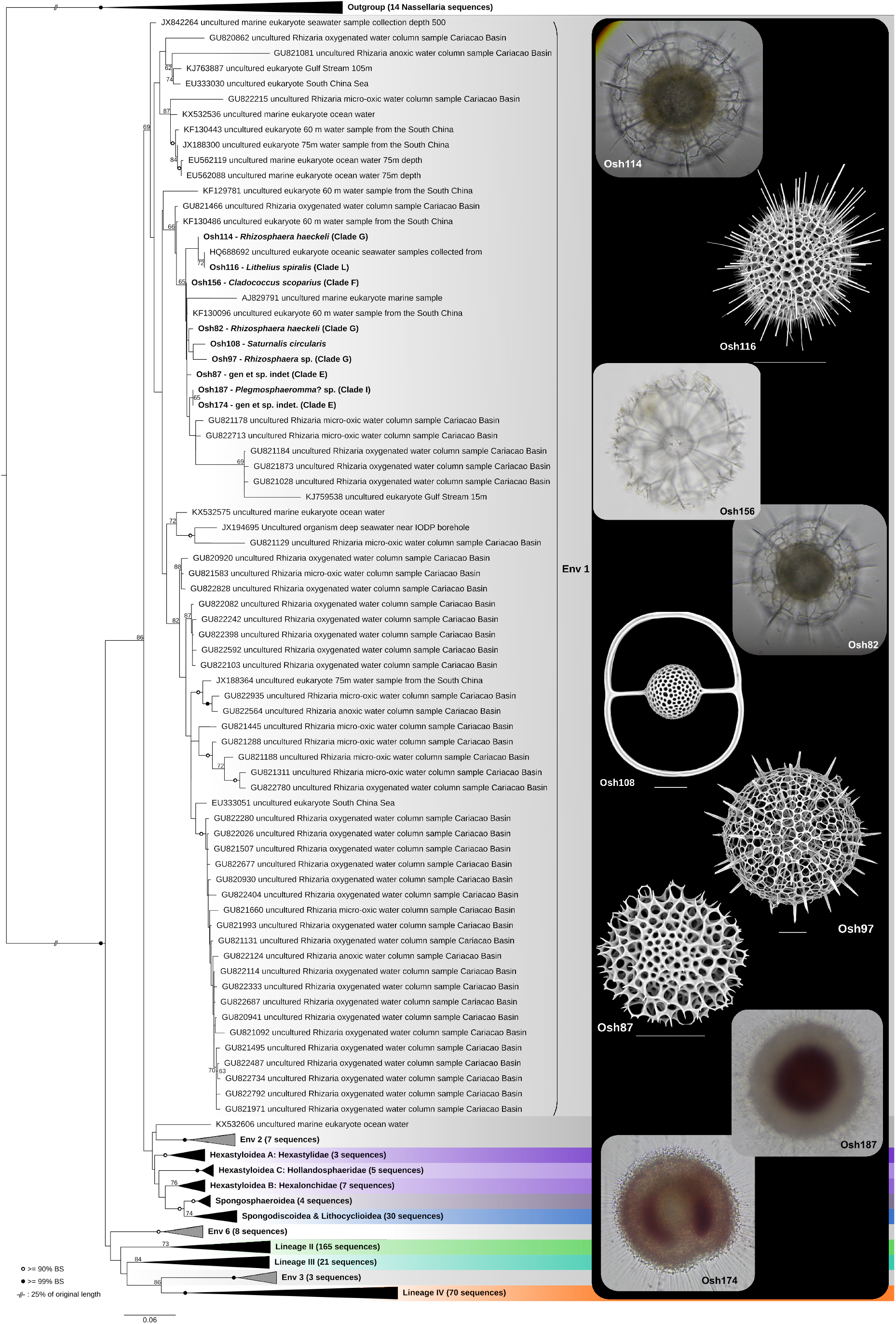
Detailed molecular phylogeny of the environmental clade Env1, showing the different morphotypes within the same phylogenetic clade. In brackets after the species name is indicated the expected clade based in the morpho-molecular framework established herein. RAxML bootstrap values (1000 replicates, BS) are shown at the nodes. Black circles indicate BS ≥ 99% and hollow circles indicates BS ≥ 90%. BS lower than 60 were removed for clarity. Branches with a double barred symbol are fourfold reduced for clarity.

To assess possible pseudogenes for clade Env1, we performed two analyses for the 18S rDNA: (i) an entropy analysis of the alignment and (ii) the GC content estimation of every sequence. No differences were found between the entropy of Env1 and that of the different lineages (Supplementary Material Fig. S4), showing an average Shannon entropy of 0.07 (±0.16 standard deviation; sd) in Env1, 0.08 (±0.19 sd) in Lineage I, 0.03 (±0.08 sd) in Lineage II, 0.11 (±0.24 sd) in Lineage III and 0.14 (±0.26 sd) in Lineage IV. The greatest entropy values correspond to the V4 hyper-variable region of the 18S rDNA gene (positions ~450-~850). Other important regions correspond to the V2 (around the position ~200), V7 (~1400) and V9 (~1720) hyper-variable regions of the 18S rDNA gene. GC content of Env1 also reveals comparable results to the other clades, where Env1 has an average GC content of 50.1% (±1.36% sd) and the average of all the sequences analyzed is of 48.9% (±1.61% sd) (Supplementary Material Fig. S5). Yet Env1 has the highest average GC content among its relatives in Lineage I and decreases towards more distal groups, finding a similar pattern in Lineage IV.

## Discussion

### Morpho-molecular classification of Spumellaria

Our study shows that the symmetry of the skeleton and the overall morphology are key features describing spumellarian phylogeny at higher rank classifications, as for other radiolarian groups such as Acantharia (Decelle et al., 2012b) or Nassellaria (Sandin et al., 2019). Central structure patterns are also well defined between clades helping to discern similar symmetric trends in distantly related clades (e.g. flat and radial symmetry in the super-families Spongodiscoidea, Spongopyloidea and Stylodictyoidea). These results contrast the recognized importance of internal structures in previous morphology-based radiolarian classifications (De Wever et al., 2001; Afanasieva et al., 2005) in which the initial spicular system was the key element for taxonomic delimitation and characterization. Yet, overall shape has always played a significant role in the further description of different taxonomic groups. We therefore establish the symmetry and the overall morphology of the skeleton of Spumellaria as main morphological markers, followed by central structure patterns at finer taxonomic ranks.

The morpho-molecular framework established herein understands the Spumellaria as polycystines radiolarians with concentric structure and a spherical or radial symmetry. This concept includes some living groups classified under the order Entactinaria in De Wever et al. (2001) such as Hexastyloidea that were already suggested to be included in Spumellaria (Yuasa et al. 2009, Ishitani et al. 2012, Matsuzaki et al. 2015). Our results also confirm the inclusion of the Entactinaria super-families Centrocuboidea and Rhizosphaeroidea within Spumellaria. The Radiolaria orders Spumellaria and Entactinaria were traditionally separated based on the absence or presence of internal structures respectively (De Wever et al., 2001). Here we show that the Entactinaria order in the sense of De Wever et al. (2001) is scattered into Clades A, B, C (from lineage I) G, H and I (from lineage III). Recent advances in imaging techniques have suggested the Spumellaria as a sub-order within the Entactinaria based on findings of homologous structures with the initial spicular system in Spumellaria among primitive radiolarian forms (Kachovich et al., 2019). Despite our results agree with Kachovich et al. (2019) in the taxonomic inclusion of both orders, we consider Entactinaria as a polyphyletic group within the Rhizaria as shown in Nakamura et al. (2020). The prevalence of the overall shape at higher taxonomic levels would therefore place the Spumellaria as the name of polycystines radiolarians with concentric structure and a spherical or radial symmetry, in contrast to Kachovich et al. (2019). The formal inclusion of most entactinarian families with extant Spumellaria provides a direct and documented link from the Cambrian to current ecosystems, improving the connection for paleo-environmental reconstruction analyses. In addition to our conclusions, there are no genetic evidences of a purely Entactinaria clade so far and at this point we can not exclude that other groups sharing these symmetric patterns might also be emended within Spumellaria, as for the Centrocubidae and Rhizosphaeridae families. Despite the demonstrated polyphyletic nature of Entactinaria, different appearance and extinction ranges along the fossil record makes it a valuable taxa for biostratigraphic studies.

Extant Spumellaria included in our study can be divided in 13 morpho-molecular clades, grouped in four main evolutionary lineages based on the combination of phylogenetic clustering support, morphological features and molecular dating. Our results confirm the monophyly of some families (e.g., Liosphaeroidea, Rhizosphaeridae, Actinommoidea or Pylonioidea) while others such as Hexastyloidea found in clades A, B and C or Spongodiscoidea previously gathering morphologies from clades E and L are polyphyletic (Matsuzaki et al., 2015). Different families included within Hexastyloidea have been classified as independent super-families in previous morphology-based taxonomic studies (De Wever et al., 2001; Afanasieva et al., 2005), as seen in the paraphyletic relationships of our phylogenetic results. For instance, flat spumellarians have been classified all together on the basis of their flat symmetry, yet the previously stated polyphyletic nature of this group is confirmed by our results, due to differences in the test and the overall morphology of the skeleton (De Wever et al., 2001). Despite our efforts, some uncertainties regarding specific families such as Heliodiscidae, Ethmosphaeridae or Saturnalidae remain yet to be described. This could be adressed by performing further single celled morpho-molecular characterization analyses on targeted specimens from a broad variety of environments (e.g. deep water). Although these families are represented by very few species and have similar symmetric and central structure patterns to other families presented herein; they could be therefore emended within one of the described clades.

### Diversity of Spumellaria

The great environmental diversity found at early diverging positions (i.e.; Env1, Env3, Env6 from Fig. 6) prevents a comprehensive exploration of the extant spumellarian morpho-molecular diversity and challenges the reliability of such molecular clades. Our analyses allowed however hypothesizing different morphologies for such environmental clades, precisely Env1. Various lines of evidences led us to conclude the existence of two different and phylogenetically distant 18S rDNA gene copies within the same specimen:

i. The notably high ratio of chimeric sequences obtained from each single cell analyzed (~13-25%) that would reflects the presence of different rDNA genes amplified with the same Spumellaria-specific primer sets from a single cell/specimen.
ii. The common relatively long branches within clade Env1 (Fig. 7) traducing the decreased accuracy and quality of sequences obtained by the sequencing of a mix of rDNA gene copies.
iii. Distantly related morpho-species unexpectedly clustered in the same phylogenetic clade Env1 (Fig 7), contrasting with the globally well supported morpho-molecular clades in Spumellaria.
iv. The presence of the same single cell specimen in 2 different and highly supported clades (Osh174 in Env1 and E2), when different amplification and sequencing runs have been carried out.

Further analyses based on the entropy and GC content of sequences clustering in Env1 showed similar trends compare to the rest of spumellarian sequences, disregarding the possibility of a pseudogene (e.g. Balakirev et al., 2014). Given the regular occurrence of symbiotic relationships within the SAR super-group (Stoecker et al., 2009, 2016; Bjorbækmo et al., 2019), the most likely explanation left for clade Env1 is the existence of a naked symbiotic group of Spumellaria developing within another, skeleton-bearing Spumellaria. The exact symbiotic nature (parasitic, mutualistic or commensalistic) of such group remains beyond the scope of this study and cannot be inferred from our data. Further imaging analysis on live or fixed specimens (e.g. fluorescence *in situ* hybridization) could help settle this hypothesis by visualizing two cells within the same skeleton or detecting skeleton-less Spumellaria. Similar associations have been already identified between benthic bathyal Foraminifera and dead tests of Xenophyophorea (Foraminifera) (Pawlowski et al., 2003; Hughes and Gooday, 2004). To our knowledge the few molecular data available for marine Heliozoa have appeared to be polyphyletic (Cavalier-Smith et al., 2015; Burki et al., 2016), as for fresh water heliozoans (Nikolaev et al., 2004). Indeed, some groups of Heliozoa, such as Gymnosphaeridae, have been moved to the Retaria lineage (precisely within Radiolaria) based on ultra-structural features of the axopodial complex (Yabuki et al., 2012). Therefore, one cannot rule out that some of these basal environmental clades are represented by heliozoan-like organisms and some of them (clade Env1) developed symbiosis within other Spumellaria.

The morpho-molecular framework of Spumellaria established in this study allows an accurate phylogenetic placement of environmental sequences for which it would not be possible to infer their taxonomic position otherwise. Most of the environmental (i.e. not morphologically described) sequences placed in our reference morpho-molecular framework are highly related to Liosphaeroidea, coming primarily from deep environments (1500-2500m) (Edgcomb et al. 2011; Lie et al. 2014). Another group displaying a large environmental diversity is the family Excentroconchidae, mainly represented by sequences coming from anoxic environments (Edgcomb et al. 2011; Kim et al., 2012). These results agree with the data obtained by morphological observations from the fossil record (De Wever et al., 2001) or from plankton and sediments traps and surface sediment materials (Boltovskoy et al., 2010; Boltovskoy and Correa, 2016), in which both the Liosphaeroidea and the Excentroconchidae are among the most abundant groups. Representatives of these families are characterized by a thin and fragile skeleton with a dense protoplasm. As previously suggested (Not et al., 2007), cell breakage may also contribute to the outstanding amount of sequences found among these taxonomic groups compared to other clades.

### Evolutionary history of Spumellaria

Our molecular clock results dated the first diversification of Spumellaria at ~515 (HPD: 659-382) Ma, in agreement with recent findings of the oldest radiolarian fossils with spherical forms, taxonomically classified in the extinct spumellarian genus *Paraantygopora* (Early Cambrian, Series 2, ca. 521-509 Ma; Zhang and Feng, 2019). This early diversification separated the two contrasting lineages I and IV, followed soon afterwards by the branching of linages II and III, during the so-called Great Ordovician Biodiversification Event (Noble and Danelian, 2004; Servais et al., 2016). At this moment many different symmetrically spherical radiolarians appeared in the fossil record (Aitchison et al., 2017; Noble et al., 2017). Yet, poor radiolarian-bearing rocks (Aitchison et al., 2017), different topologies of the different marker genes and the short phylogenetic distance at these nodes in the concatenated phylogenetic analysis make relationships between lineages obscure. From this period until the end of the Paleozoic the diversification of extant lineages is low and basically restricted to the end of the Carboniferous (358.9 – 298.9 Ma) when lineages III and IV start diversifying.

All over the Paleozoic, extinct specimens previously attributed to Entactinaria show a great diversity that mostly became extinct towards the beginning of the Mesozoic (De Wever et al., 2003, 2006). Some representatives of this group have been tentatively considered as Nassellaria in latest classifications and later suggested as ancestors of this order due to overall morphology and molecular clock results (Noble et al., 2017, Sandin et al., 2019). The morpho-molecular framework here proposed allows the attribution of some radiolarian extinct groups with spherical or radial symmetry as possible ancestors of the extant Spumellaria, that can be reflected in the long branches of clades belonging to lineages II, III and IV (Fig. 1). These phylogenetic patterns may be interpreted as a bottleneck effect, in which many groups got extinct before radiation.

At the beginning of the Mesozoic the first extant representatives appear with the diversification of Lineage II (Liosphaeroidea) and Hexastyloidea. Although, the polyphyletic nature of Hexastyloidea shows a later diversification within the clades (A, B and C), explaining why the Triassic (251.9 – 201.3 Ma) genera does not resemble that of the Cenozoic (66 – 0 Ma) (O’Dogherty et al., 2011). During this period, Lineage III also diversifies and all clades from lineage IV are already separated from each other, yet their diversifications are not happening until the end of the Mesozoic. Similar diversification patterns at the beginning of the Mesozoic have been reported in living Nassellaria inferred from the combination of the molecular clock and the direct observation of the fossil record (De Wever et al., 2006; Sandin et al., 2019); in which ancient forms, presumably extinct during the end-Permian extinction (Sepkoski, 1981; De Wever et al., 2006; Takahashi et al., 2009; O’Dogherty et al., 2011; Aitchison et al., 2017) led to the diversification of living groups.

The second part of the Mesozoic is characterized by a stepped diversification of Spumellaria, as already described for Nassellaria (Sandin et al., 2019) and Foraminifera (Leckie et al., 2002; Hart et al., 2003). This phenomenon are classically explained by the onset of a global oceanic anoxia during the Jurassic (Jenkyns, 1998) and a series of Oceanic Anoxic Events during the Cretaceous (Schalanger and Jenkyns, 1976; Erbacher et al., 1996; Jenkyns, 2010; Yilmaz et al., 2012; Kemp and Izumi, 2014). In contrast to Nassellaria diversification, towards the end of the Jurassic (201.4 – 145 Ma) and beginning of Cretaceous (145 – 66 Ma), there is a big increase in Spumellaria diversification where most of the clades suddenly diversified, also reported in the fossil record (Kiessling, 2002; De Wever et al., 2003). Such differences between diversification patterns of Spumellaria regarding other radiolarian groups is a question already rose in recent studies (Kachovich et al., 2019). Both extant Nassellaria and Spumellaria share similar environmental preferences (Suzuki and Not, 2015). Yet Spumellaria tend to possess a larger protoplasmic volume (Takahashi, 1982; Des Combes and Abelmann, 2009). There is also a drastic diversification of flat spumellarians, a features that increases the surface/volume ratio of a given organisms. Probably such morphological adaptations provided an advantage to thrive during low nutrient availability periods, as known to be case during the Jurassic (Cárdenas and Harries, 2010). Once the conditions became more favorable heterotrophic Spumellaria started a sustained diversification that followed all over the Cenozoic (66 – 0 Ma). Similar diversification patterns linked to climatic oscillations over the Cenozoic were found in Nassellaria (Sandin et al., 2019), Foraminifera (Hallock et al., 1991) and coccolithophores (Bown et al., 2004).

## Acknowledgments

This work was supported by the IMPEKAB ANR 15-CE02-0011 grant and the Brittany Region ARED C16 1520A01, the Japan Society for Promotion of Science KAKENHI Grand No. K16K0-74750 for N. Suzuki and “the Cooperative Research Project with the Japan Science and Technology Agency (JST) and Centre National de la Recherche Scientifique (CNRS, France) “Morpho-molecular Diversity Assessment of Ecologically, Evolutionary, and Geologically Relevant Marine Plankton (Radiolaria)”. We are grateful to the CNRS-Sorbonne University ABiMS bioinformatics platform (http://abims.sb-roscoff.fr) for providing computational resources. The authors are grateful to the MOOSE observation national network (funded by CNRS-INSU and Research Infrastructure ILICO) which sustain the annual ship-based hydrographic sections in the northwestern Mediterranean Sea (MOOSE-GE), as well as John Dolan for hosting us multiple times at the Laboratoire d’Océanographie of Villefranche sur Mer. We are greatly thankful to Cedric Berney for the phylogenetic advises and the valuable help on the interpretation of the “symbiotic” clade, as well as Vasily Zlatogursky for his contributions and feedback on the heliozoan-like organism.

## Supplementary Data

**Table S1.** List of specimens used to obtain spumellarian phylogeny.

**Table S2.** List of morphological characters (traits) and their states (1 to 3) used for the ancestral state reconstruction analysis.

**Table S3.** List of environmental sequences related to Spumellaria.

**Figure S1.** A compared molecular phylogeny of Spumellaria inferred from the 18S (on the left) and partial 28S (D1-D2 regions, on the right) rDNA genes. Trees were obtained by using a phylogenetic Maximum likelihood method implemented using the GTR + γ + I model of sequence evolution. PhyML bootstrap values (100 000 replicates, BS) are shown at the nodes (BS). Black circles indicate BS ≥ 99%. Hollow circles indicate BS ≥ 90%. Branches with a double barred symbol are fourfold reduced for clarity.

**Figure S2.** Light Microscopy (LM) and Scanning Electron Microscopy (SEM, when available) images of live Spumellaria specimens used in this study for phylogenetic analysis in Fig. 1. Scale bars (when available) = 50μm.

**Figure S3.** Parsimony Ancestral State Reconstruction analysis based in the resulting phylogenetic tree for the 6 characters chosen. Relevant nodes are increased for clarity.

**Figure S4.** Shannon entropy of every position of the 18S rDNA gene alignment used to obtain phylogenetic tree of figure 7 for the different lineages (excluding Env 1 from Lineage I) and for Env 1 independently. Lines correspond to the smoothed generalized linear model with the 99% confidence interval under the grey area.

**Figure S5.** Boxplot showing the median (thick horizontal lines), the interquartile range (box), the minimum and maximum values (1.5 coefficients; vertical lines) and potential outliers (dots) of the GC content of every sequence used in this study grouped by its taxonomic affiliation showed in figure 6 & 7.

