## Supplementary material for "Morpho-molecular diversity and evolutionary analyses suggest hidden life styles in Spumellaria (Radiolaria)": TableS2.pdf

**Table S2.** List of morphological characters (traits) and their states (1 to 5) used for the ancestral state reconstruction analysis.

| | | Traits | | | | | |
|---|---|---|---|---|---|---|---|
| | | Skeleton shape | Symmetry | Central structure | Internal cavity | Number of spines arisen from central structure | Number of distinctive cortical shell |
| State | 0 | Spherical | Line[b] | Empty | Wide space | 0 | 1 |
| | 1 | Spherical modified[a] | Spherical or plane | Entactinaria-type, fused fibers or soft spongy[e] | Space dividers | 6 | 2 |
| | 2 | Cubic-, box-, dice-shaped | Reflection and/or plane[c] | Double medullary shell or highly dense centre[f] | Dense[h] | More or less than 6 | 3 |
| | 3 | Flat | Circular or rotational[d] | Complex[g] | Dense with pylome[i] | Significantly more than 6 | More than 3 |

[a] Cylindrical or spherical to ellipsoidal.

[b] Line symmetry for one, three or *n* axes.

[c] Reflection symmetry for one axes and plane symmetry or reflection symmetry for three axes.

[d] Circular, tria-rotational, tetra-rotational or n-fold rotational and plane symmetry.

[e] Tetrapetaloid, cube or Rhizosphaerid-type microsphere, fibers fused at the centre or very delicate framed central structure.

[f] Spherical or flatten double medullary shell, spherical microsphere with decussate 4 radial beams, margarita-type microsphere or spinose and homogeneous dense structure.

[g] Pit-like microsphere with tunnel-like pylome, spinose with straight radial beams, tetrapyd system or phorticid system microsphere.

[h] Dense to coarse homogeneous skeletons or dense concentric structure with or without a non-walled pylome.

[i] Dense concentric structure with a tunnel-like pylome.
