## Supplementary figures and images for "Morpho-molecular diversity and evolutionary analyses suggest hidden life styles in Spumellaria (Radiolaria)"

### FigS1.pdf

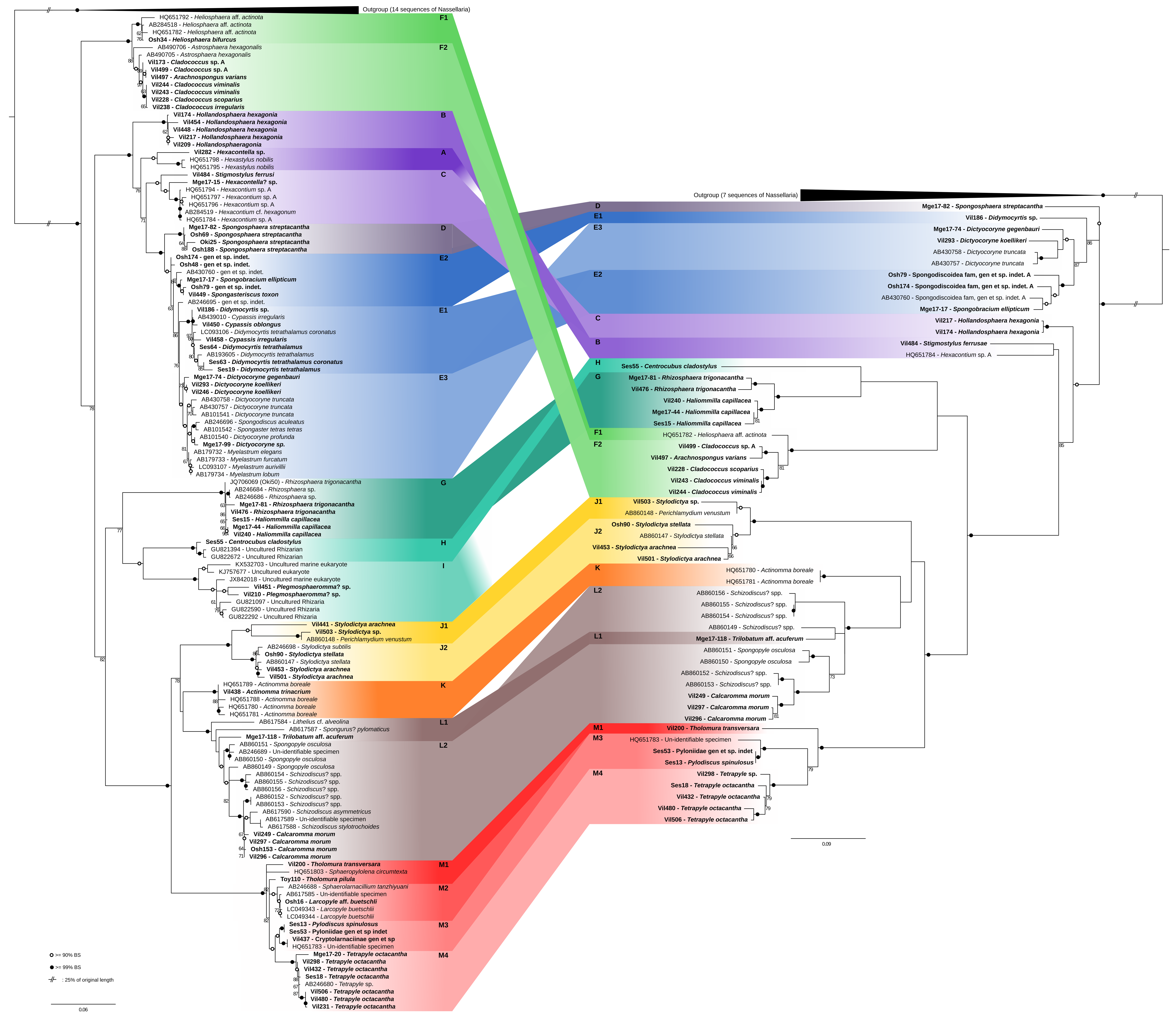

### FigS2.pdf

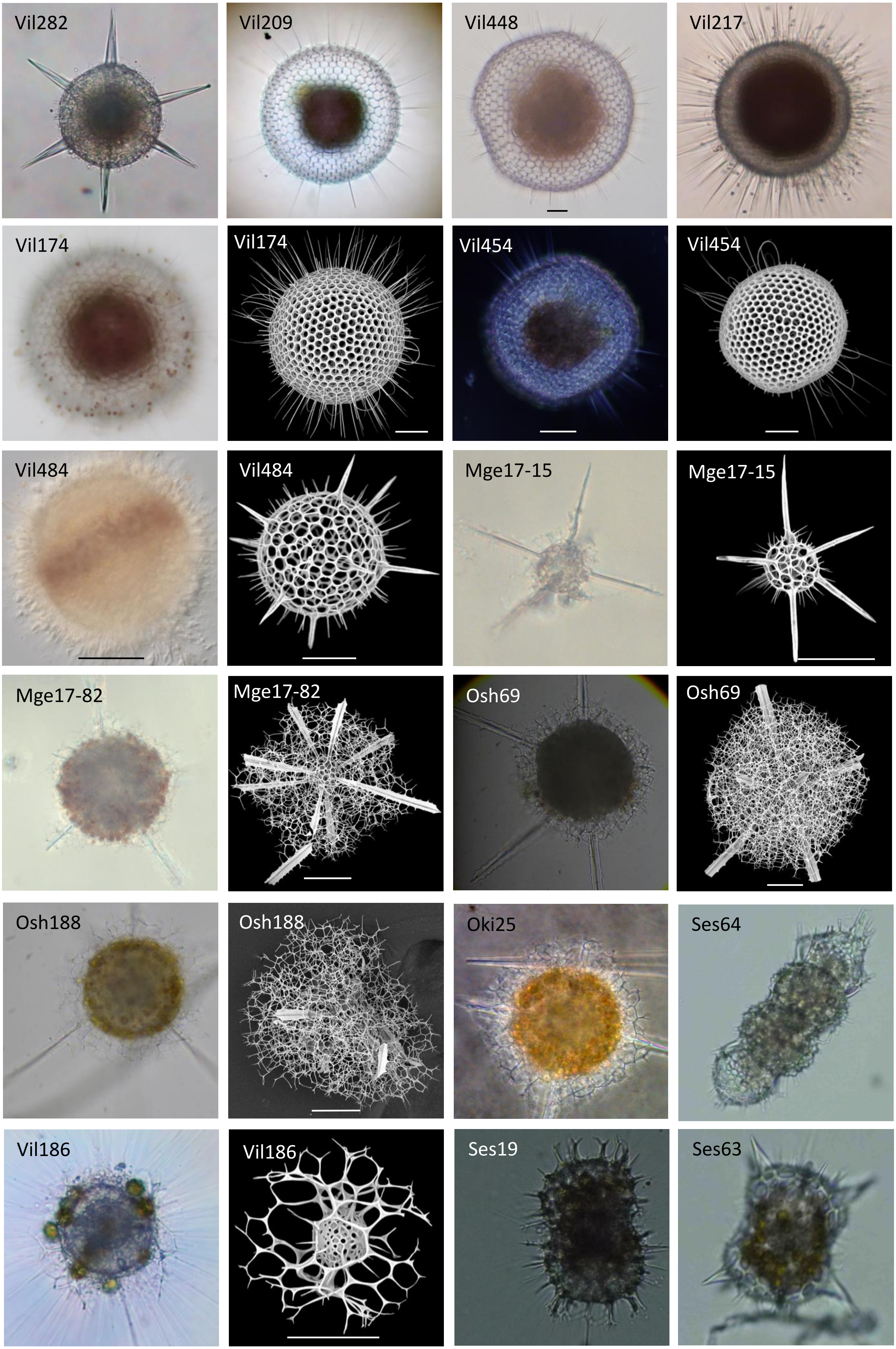

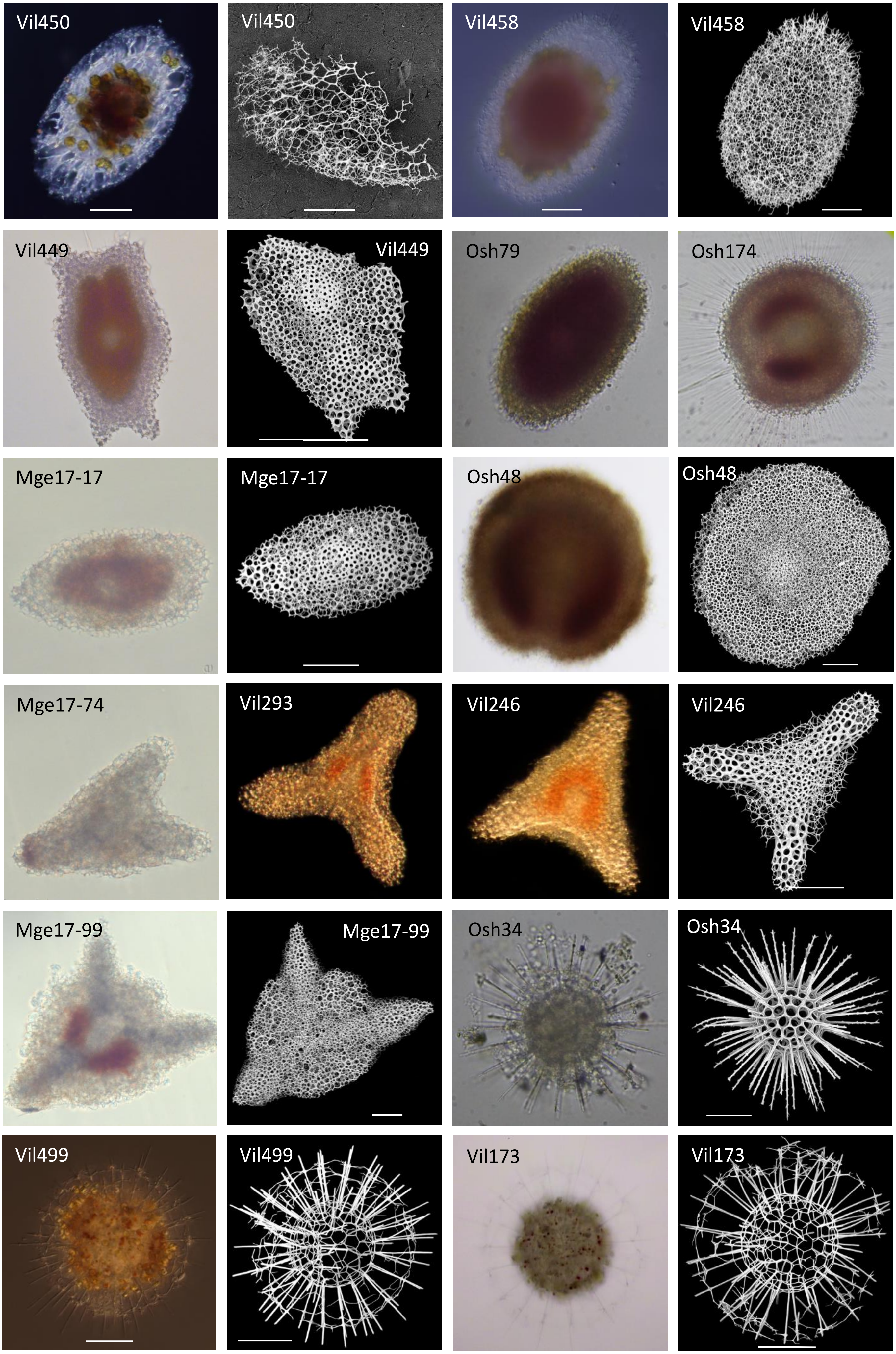

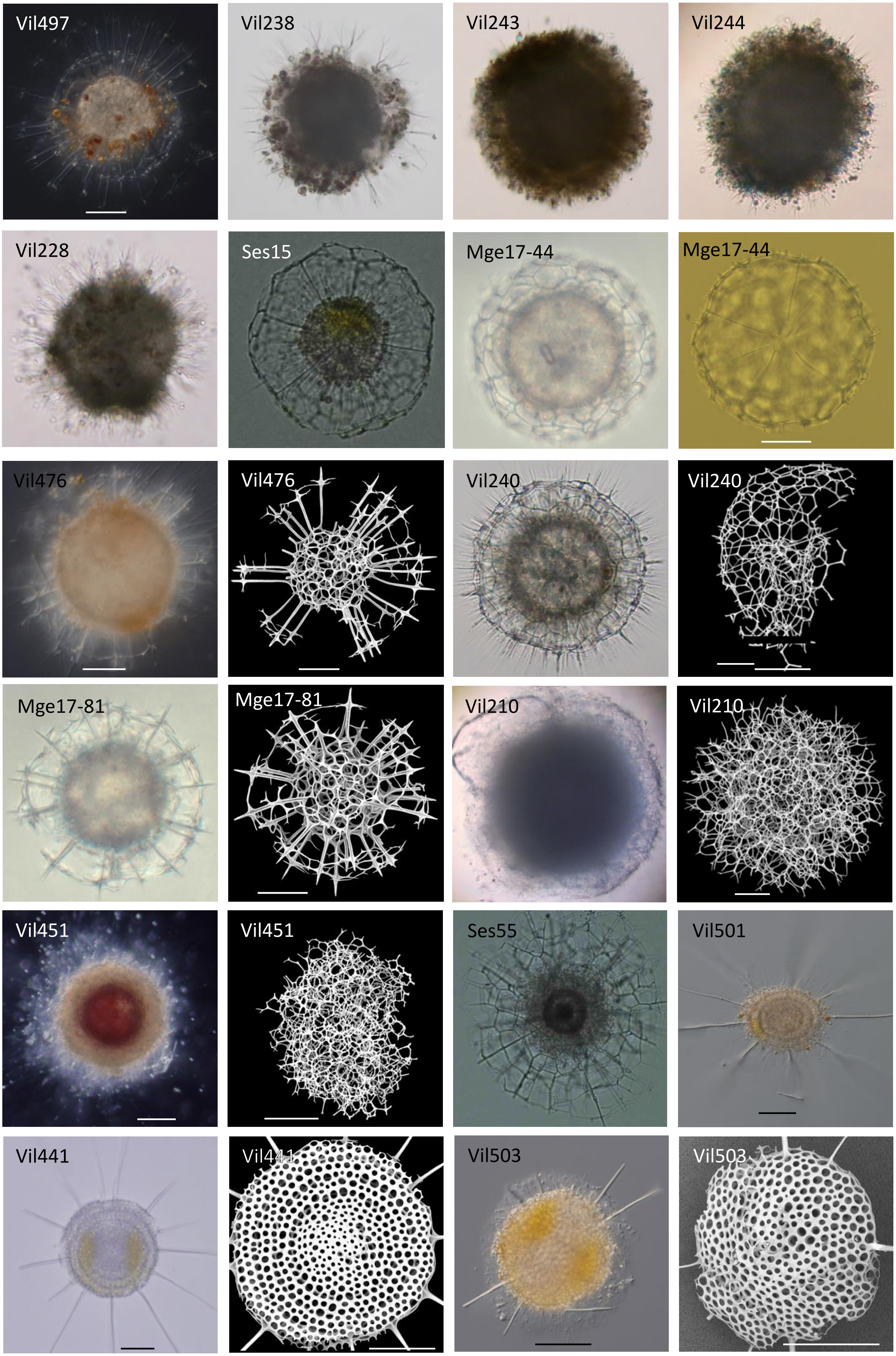

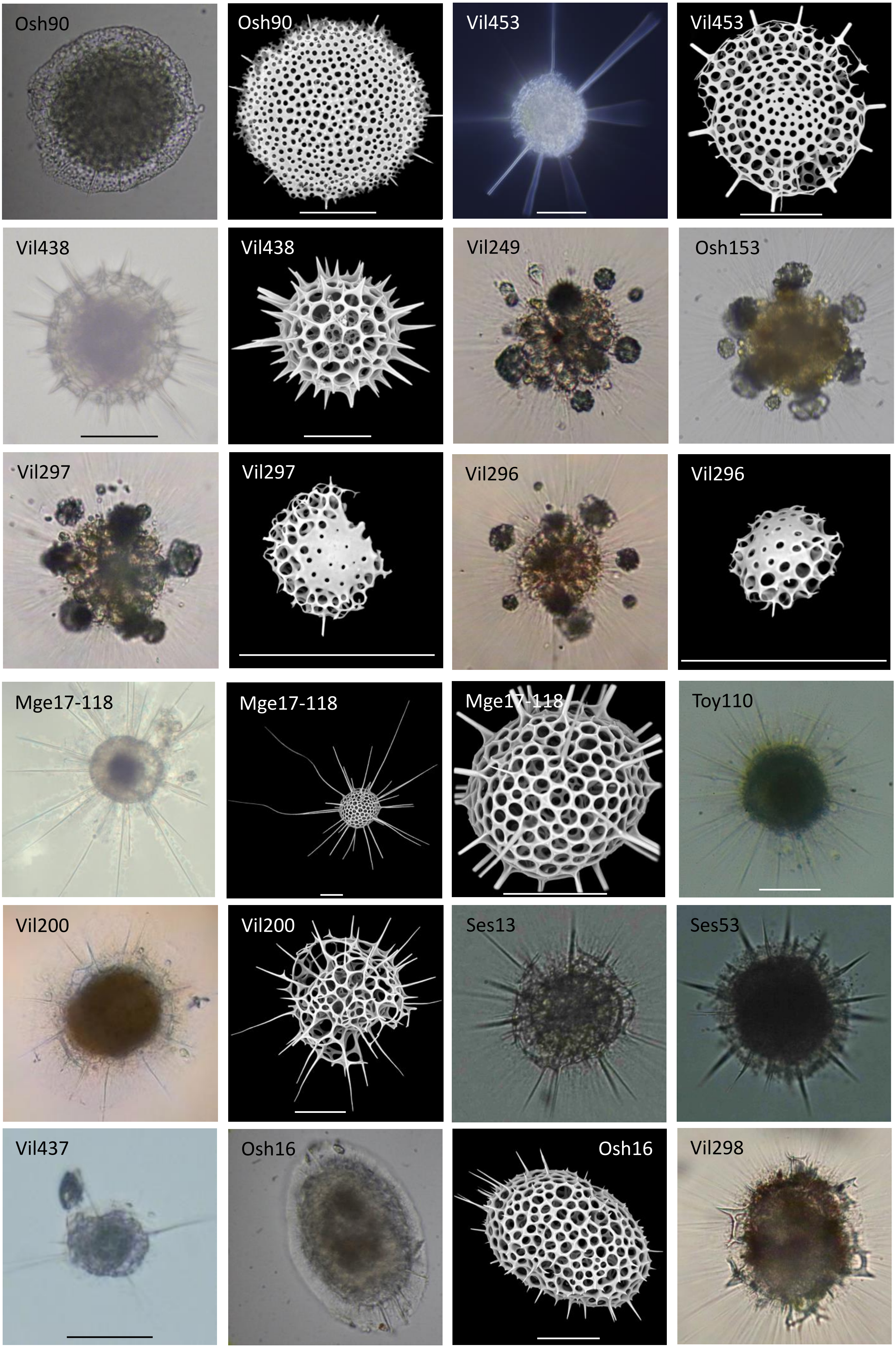

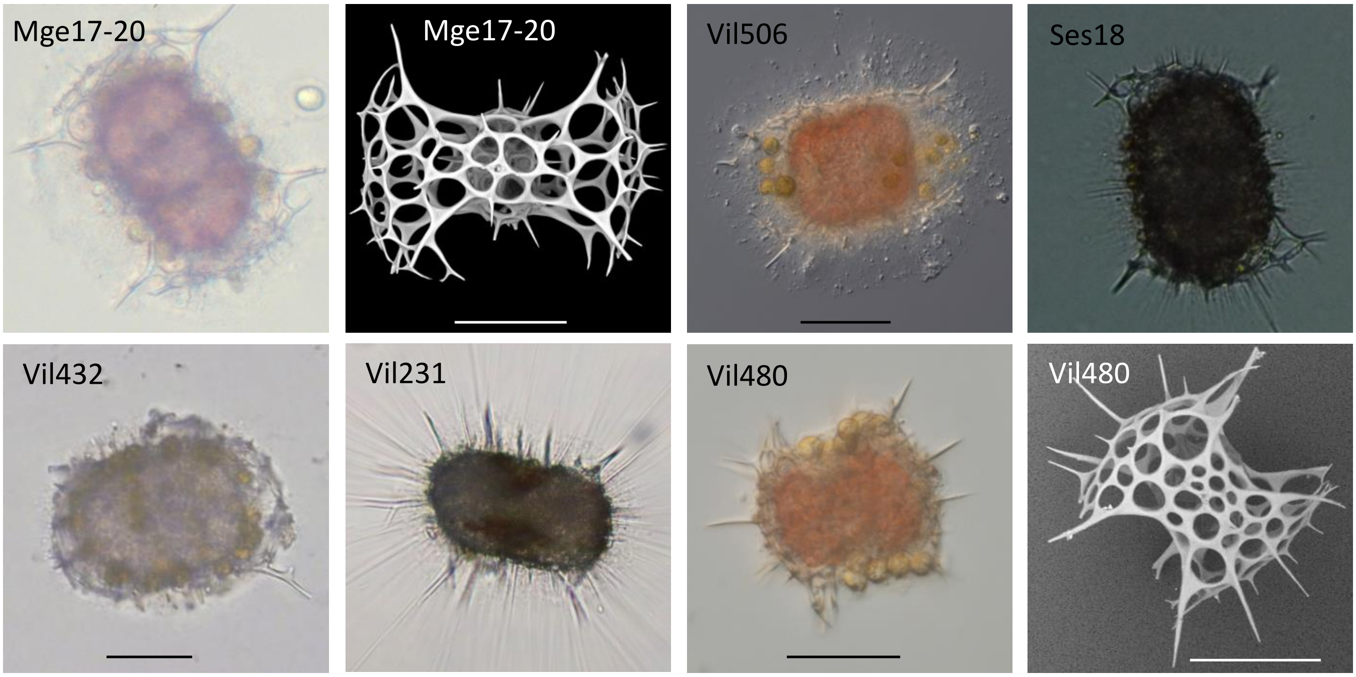

### FigS3.pdf

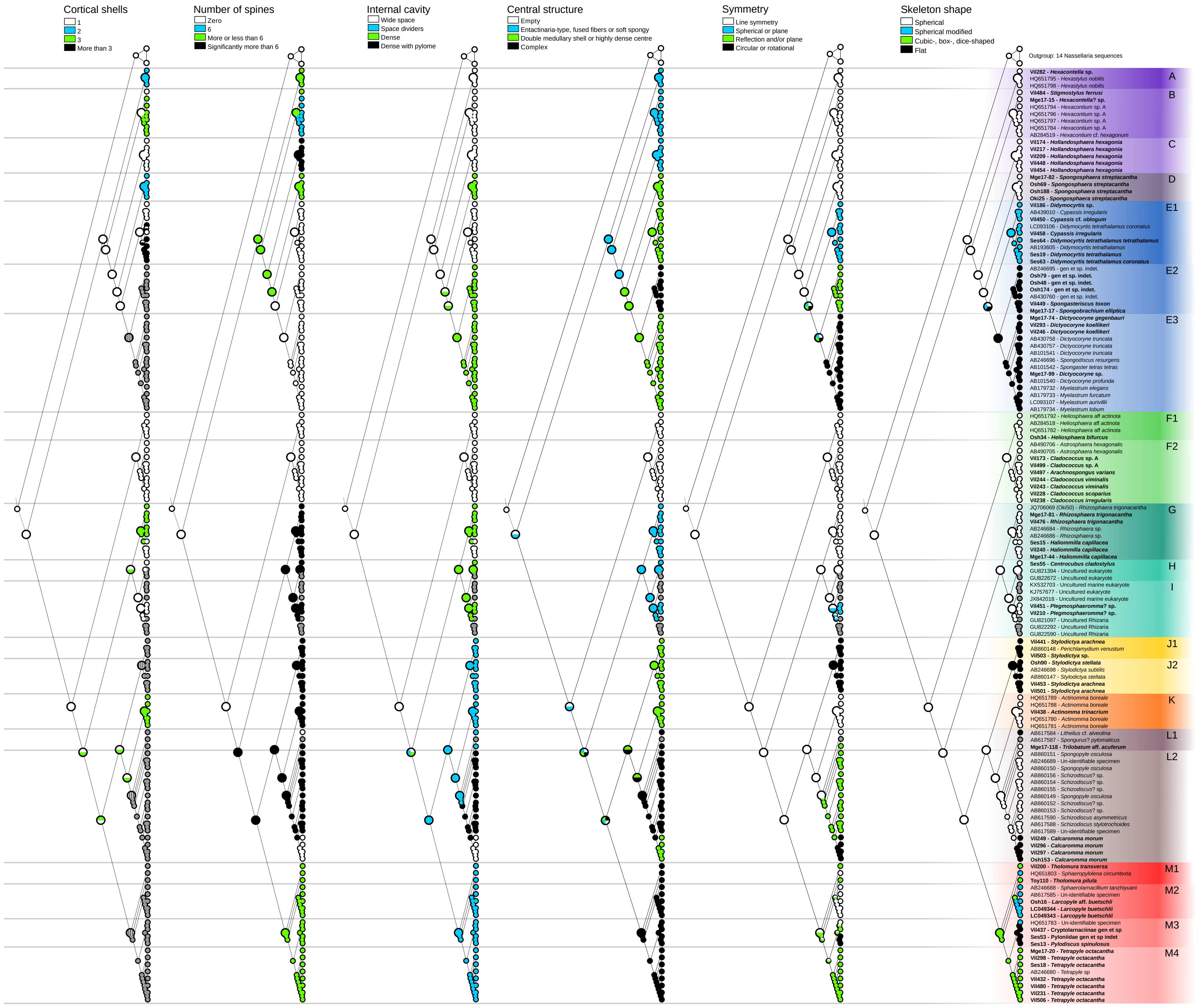

### FigS4.pdf

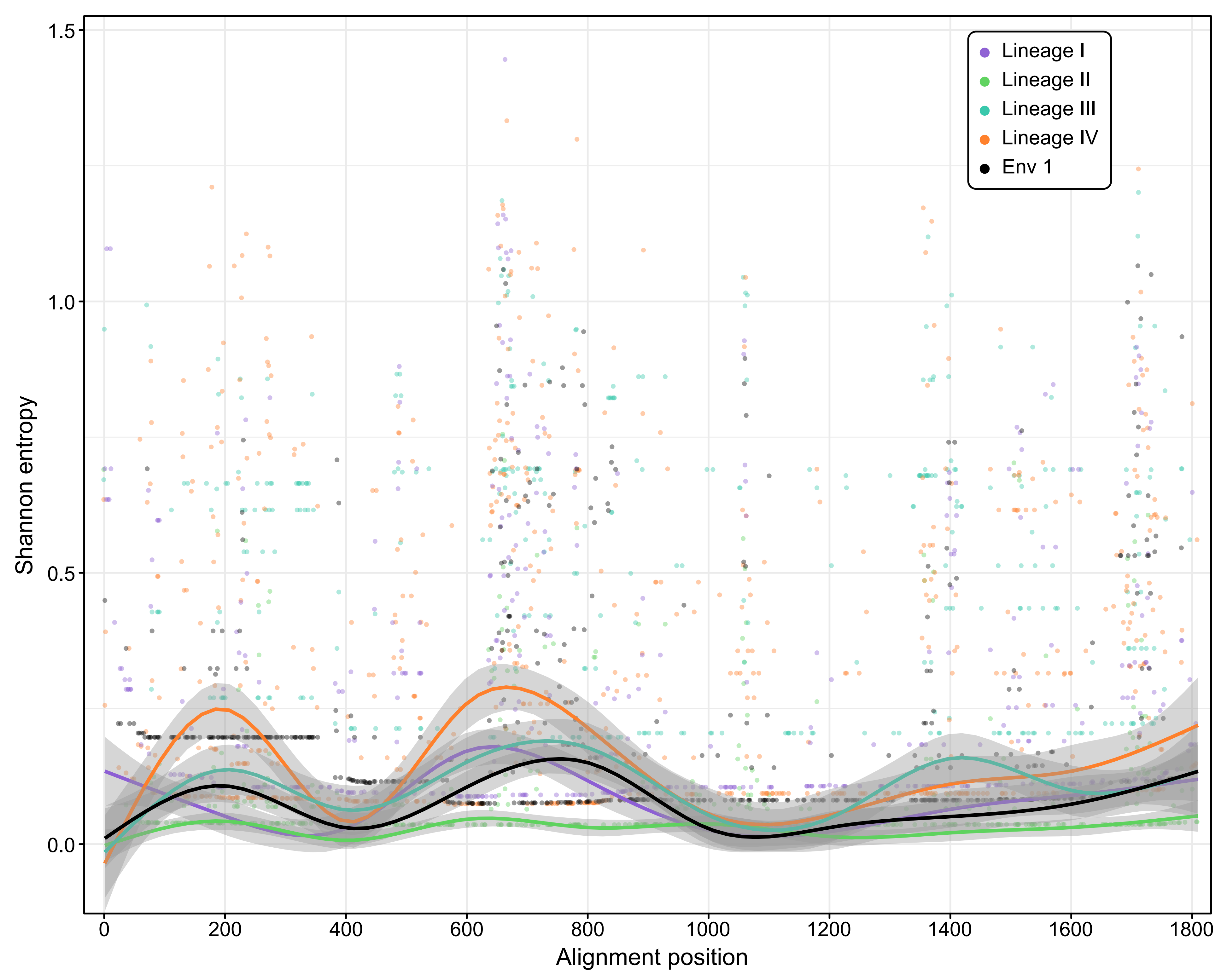

### FigS5.pdf

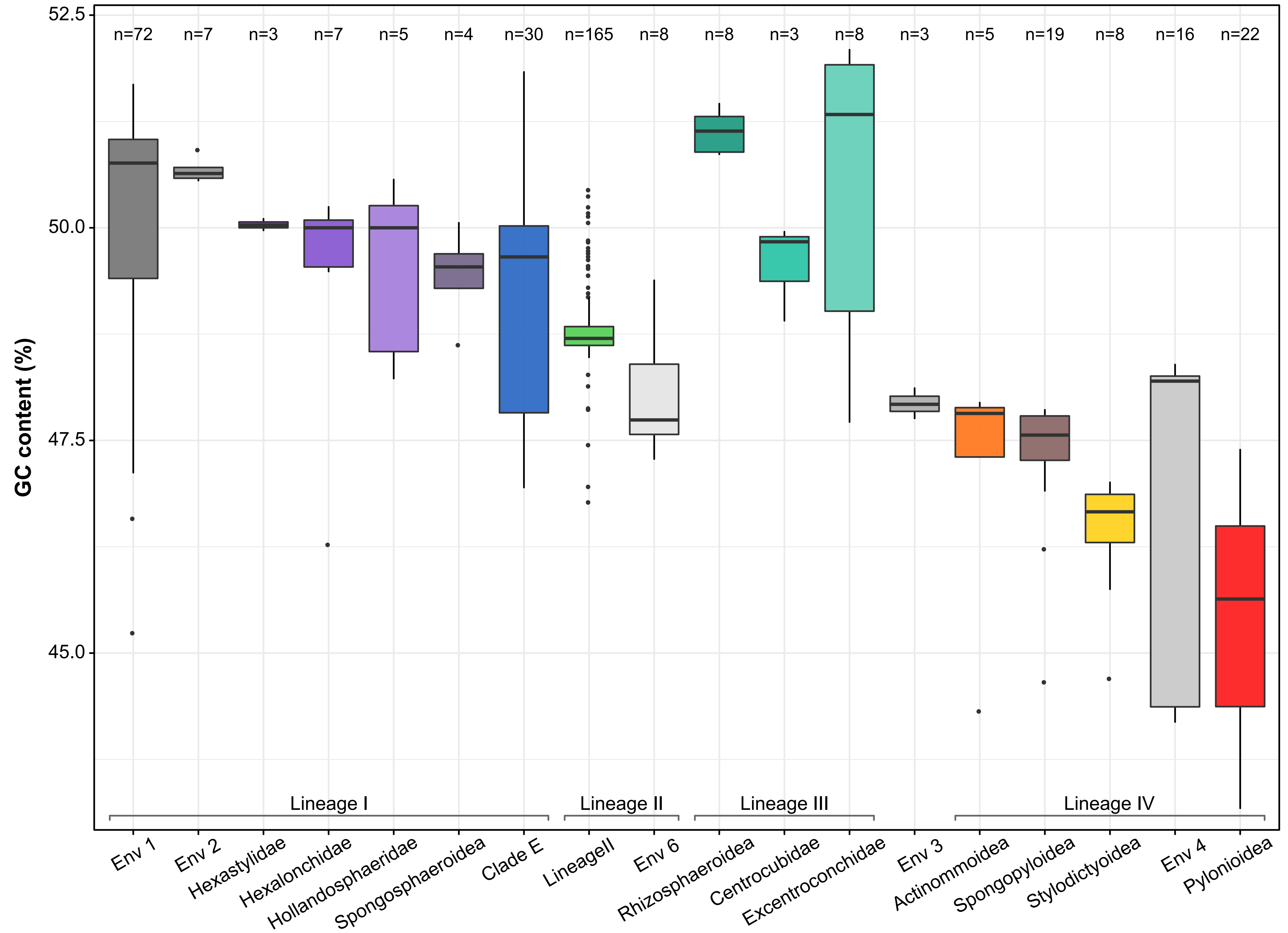
